# Palmitate-induced mitochondrial damage restricts histone acetylation in CD8^+^ T cells to impair anti-tumor immunity

**DOI:** 10.1101/2025.01.09.631811

**Authors:** Silvia Tiberti, Martina Romeo, Letizia Rumiano, Carina B Nava Lauson, Ilir Sheraj, Carlotta Catozzi, Roberta Noberini, Davide Olivari, Tiziano Dallavilla, Serena Galie’, Mattia Ballerini, Alessia Loffreda, Martin H. Schaefer, Tiziana Bonaldi, Luigi Nezi, Sara Sdelci, Teresa Manzo

## Abstract

Accumulation of lipids in the tumor microenvironment (TME) is a feature of several solid tumors and increased palmitate (PA) availability fosters tumor progression and metastases. The intrinsic effects of PA on cancer cells are well understood, but its role in modulating CD8^+^ T cells (CTL) functional performances remains elusive. Here, we found that PA alters the mitochondrial metabolism of CTL and prevents their effector functions in an irreversible manner, resulting in impaired antitumoral immunity. Mechanistically, PA-induced mitochondrial block demotes histone acetylation and chromatin accessibility and decrease transcription of genes promoting DNA replication and production of effector molecules. We identified the metabolic enzyme Sphingosine Kinase 2 (SPHK2) as a molecular target of PA in establishing CTL dysfunction. Consistently, pharmacological inhibition of SPHK2 restored CTL mitochondrial fitness, effector functions and anti-tumor potential. Thus, we reveal a critical function of PA in tumor progression by undermining CTL antitumor immunity and highlight the therapeutic potential of inhibiting SPHK2 activity to optimize T cell functionality.

## INTRODUCTION

CD8^+^ T cell (CTL) functions are tightly dependent on the metabolic landscape in which they operate (*1*), where nutrient availability has certainly a crucial role in shaping their cellular metabolism and behavior (*2–5*).

Lipids serve as “metabolic messengers” essential for energy production, membrane structure and signal transduction; however, an excess of lipids has been functionally linked with tumor progression (*6–8*). Beyond their recognized cancer intrinsic effects, lipids are emerging as key driver of T cell activity (*9*, *10*) and accumulation of specific lipids within the TME has been set as a common mechanism of immune-evasion in different types of tumors (*11–16*), thus establishing a link between lipid metabolism, metabolic fitness and immunosurveillance. However, whether tumor-derived lipids are detrimental to T cell functionality and how they regulate different CTL fate decisions has not been thoroughly investigated yet.

The long-chain fatty acid palmitic acid (PA) is produced by cancers cells and accumulates in the surrounding TME, leading to accumulating alterations in the epigenetic landscape of tumor cells that foster oncogenic transformation and dissemination (*17*, *18*). Despite our understanding of the cancer cell–intrinsic regulation by PA, the tumor cell–nonautonomous effects of PA in the TME still remain poorly understood. Here, we set to dissect whether PA can directly affect anti-tumor immunity focusing on CTL, which play a main role in the anticancer response.

We reported PA as a major negative regulator of CTL by imprinting a metabolic dysfunctional state associated with epigenetic perturbations, which dampens proper activation in CTL and, consequently, hampers their antitumoral capabilities. Thus, beyond its known pro-tumoral mechanism, we discovered a new tumor-extrinsic role for PA in promoting oncogenic transformation by impeding CTL anti-tumor immunity with potential therapeutic applications to enhance CTL function and immunotherapy efficacy.

## RESULTS

### Palmitate restricts CTL activation and functionality

Palmitate (PA) exerts an important tumor-intrinsic function by activating a pro-metastatic signaling that promotes tumor progression (*17*, *18*). In addition, it has also been reported to be characteristic of intrapancreatic dysfunctional CTL (*12*), implying a tumor-extrinsic effect of PA in the lipid-rich TME.

However, whether and how PA affects CTL responsiveness remains unclear. To elucidate the role of PA on CTL activation and function, we established an *in vitro* model in which CTL are activated in the presence or absence of PA and downstream effector functions were assessed (Figure 1A). After CTL activation in the presence of PA, blast and cluster formation were decreased (Figure 1B and S1A), in line with a marked reduction of S6 ribosomal protein (S6) phosphorylation (Figure S1B), the downstream target of mTORC1, which controls cell size and biogenesis (*19*, *20*). Thus, PA reduces blastogenesis in CTL, a phenomenon associated with abrogated CTL activation (*21*). Given these observations, we investigated the effects of PA on early CTL activation. Upon PA exposure, CTL hold a defect in IL2 production - as determined by decreased IL2 intracellular staining (Figure 1C), as well as lower surface expression of typical activation markers, such as CD44 and CD69 (Figure S1C), coupled with a significantly reduced proliferation (Figure 1D). Of note, PA retained the ability to limit activation of optimally activated stimulated CTL (Figure S1D-E). Altogether, these results demonstrate that PA counteracts T cell activation.

**Figure 1.**
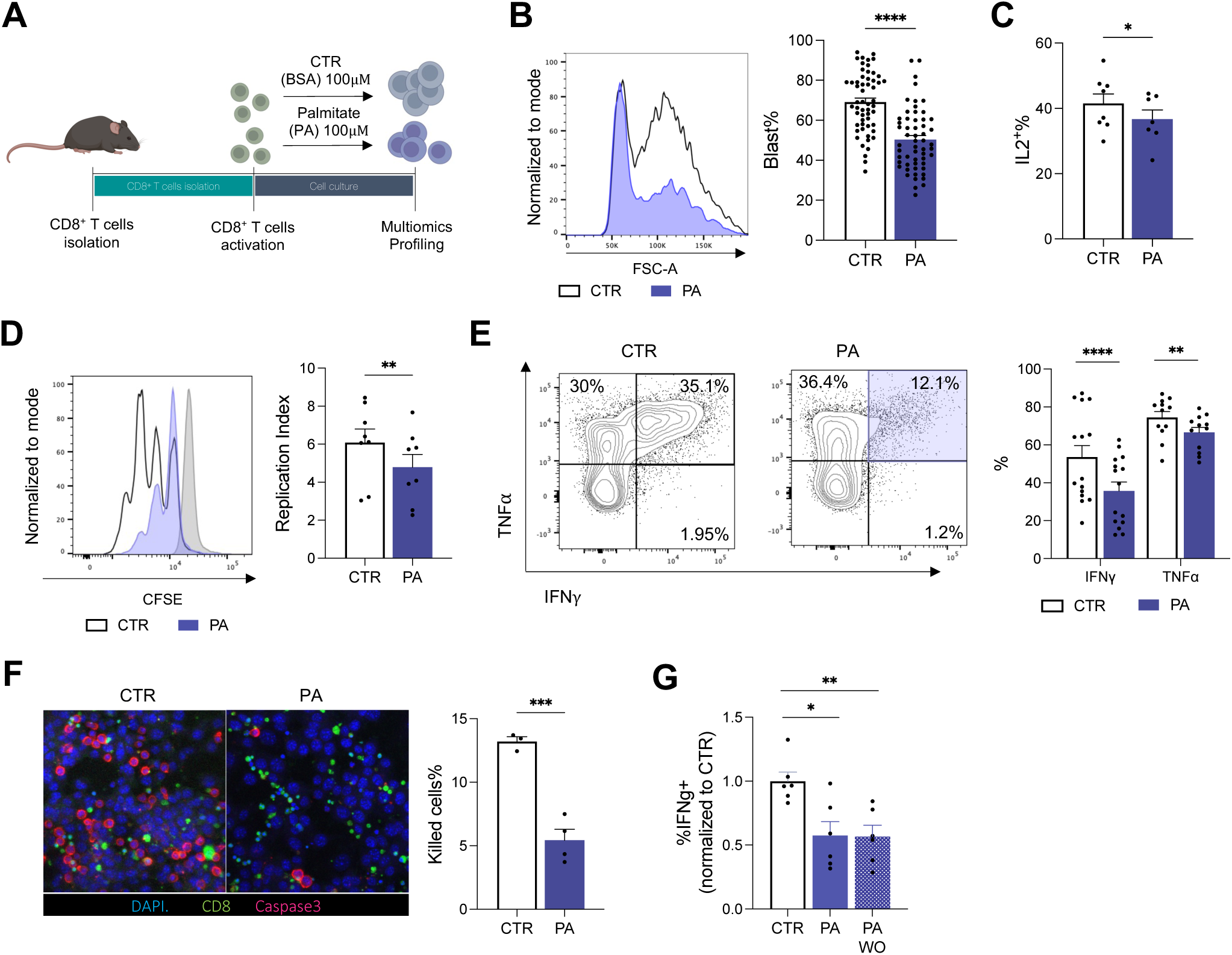
Palmitate dampens CTL functionality. **(A).** Experimental design. CTL isolated from C57.BL6 mice and activated with anti-CD28, anti-CD3 and IL2 in presence or absence of palmitic acid (PA). **(B).** FSC-A representative histogram and blast quantification (N=57). **(C).** Quantification of IL2 production (N=12). **(D).** Representative Carboxyfluorescein succinimidyl ester (CFSE) profile and relative replication index of PA- and CTR-CTL after 72h of culture. Naïve CTL were used as control (N=8). **(E).** Representative counter plot and quantification of IFNγ and TNFα production (N=12). (**F).** Immunofluorescence imagines of killing assay and relative quantification of Casp3/DAPI after 24h of coculture of OTI±PA and B16OVA in 1:1 ratio. (N=3). CD8 in purple, Caspase3 (Cas3) in green and DAPI in Blue. **(G).** Quantification of IFNγ production after 48h of culture followed by 48h with PA (PA) or wash out of PA (PA WO) compared with untreated cells (CTR). (N=6).

Since the processes of CTL proliferation, activation and cellular differentiation are tightly interconnected, we reasoned that PA might also affect CTL effector functions. PA restrains the ability of CTL to secrete effector cytokines - as demonstrated by a significant decrease in the IFNg and TNFa (Figure 1E) - alongside with a significant reduction in the expression level of Blimp1 (Figure S1F), a transcription factor necessary for the development of the effector function of CTL (*22*), further supporting a defect in the ability of PA-CTL to properly differentiate into effector CTL. Finally, to assess whether the observed defects had functional consequences, we measured the anti-tumor capabilities of CTR- and PA-CTL by measuring the antigen-specific killing of B16 ovalbumin-positive tumor cells (B16-OVA) by OVA-specific TCR transgenic T cells (OVA-CTL) that were activated in the presence of PA or left untreated. PA-CTL displayed a lower antigen-specific tumor killing capability of PA-CTL in comparison with CTR-CTL, as highlighted by the lower frequencies of Caspase3^+^ tumor cells (Figure 1F). Importantly, removing PA from the culture medium did not alleviate the observed CTL defects (Figure 1G), demonstrating that the effect of PA on CTL is not reversible. Likewise, functionality cannot be restored in tumor-infiltrating CTL exposed to a lipid-rich TME (Figure S1G-H), in which we previously demonstrated PA accumulation (*12*). Thus, we conclude that PA leads to irreversible global defects in CTL activation and effector functions, impairing their antitumor killing.

### Palmitate severely perturbs CTL mitochondrial metabolism

Next, we reasoned that PA instructs commitment to T cell dysfunction by disturbing mitochondrial fitness - a feature responsible for decreased effector functions and increased exhaustion in CTL. To this aim, we studied how PA impact on CTL mitochondrial function by utilizing different metabolic assays.

First, we used multiparametric Flow Cytometry to directly determine *ex vivo* physical parameters of mitochondria at single cell level using MitoTracker Orange and MitoTracker Green FM, which are indicative of mitochondrial function and mass, respectively. PA-CTL showed decreased mitochondrial functionality - detected as a drop in the mitochondrial potential with MitoTracker Orange staining (Figure S2A) - together with a decreased mitochondrial mass (as assessed by MitoTracker Green FM) (Figure S2B). Next, we applied immunofluorescence to detect mitochondrial morphology, which has been shown to shape metabolic reprogramming during T cell activation (*23*) and observed that, compared to CTR-CTL, PA-CTL showed significantly decreased activity and size of functional mitochondria (Figure 2A-C). This was further supported by electron microscopy analysis that highlighted the presence of smaller mitochondria in PA conditions (Figure 2D). To independently confirm the data, we measured the metabolic flux of CTL upon PA treatment. When compared to CTR-CTL, PA-CTL are characterized by a lower bioenergetic profile (Figure 2E), and lower oxygen consumption rate (OCR) (Figure 2F) together with a compromised glycolytic function (Figure S2C), spare respiratory capacity (Figure 2G) and ATP production (Figure 2H). Strikingly, all the electron transport chain (ETC) complexes were substantially reduced in PA-CTL (Figure S2D-E), indicating that PA might interfere with cellular metabolism by disturbing mitochondrial respiration at the level of ETC. Thus, PA treatment impairs mitochondrial metabolism in CTL. Finally, to address whether mitochondrial dysfunction precedes or follows impaired CTL activity, we performed a kinetic experiment dissecting the dynamic of functional effects induced by PA on CTL. Remarkably, PA initially impacted metabolic fitness by inducing a significant drop of mitochondrial activity at 24 hours (Figure S2F), followed by a compromised activation and differentiation into effector CTL after 48 hours of treatment (Figure S2G-I). Thus, our data support a model in which PA, by disrupting mitochondrial fitness, impairs CTL functional activity. Alongside the ability of lipids to mediate a context-dependent regulation of cellular processes, when tested on different tumor cell lines, PA was able to raise their mitochondrial activity (Figure S2J) and bioenergetic profile (Figure S2K-L), which might contribute to explain the reported pro-tumoral effect of PA (*17*, *18*). Consequently, by testing the ability of PA to favor tumor formation, we reported that B16 melanoma tumor cell line pre-treated *in vitro* with PA grew significantly more compared to control untreated B16 cells (Figure S2M). Combined these data provide an additional mechanism by which PA might foster tumor progression and metastasis by enabling increased tumor metabolic fitness on one hand and disrupting mitochondrial and functional fitness in CTL on the other hand, thus dampening their anti-tumor efficacy.

**Figure 2.**
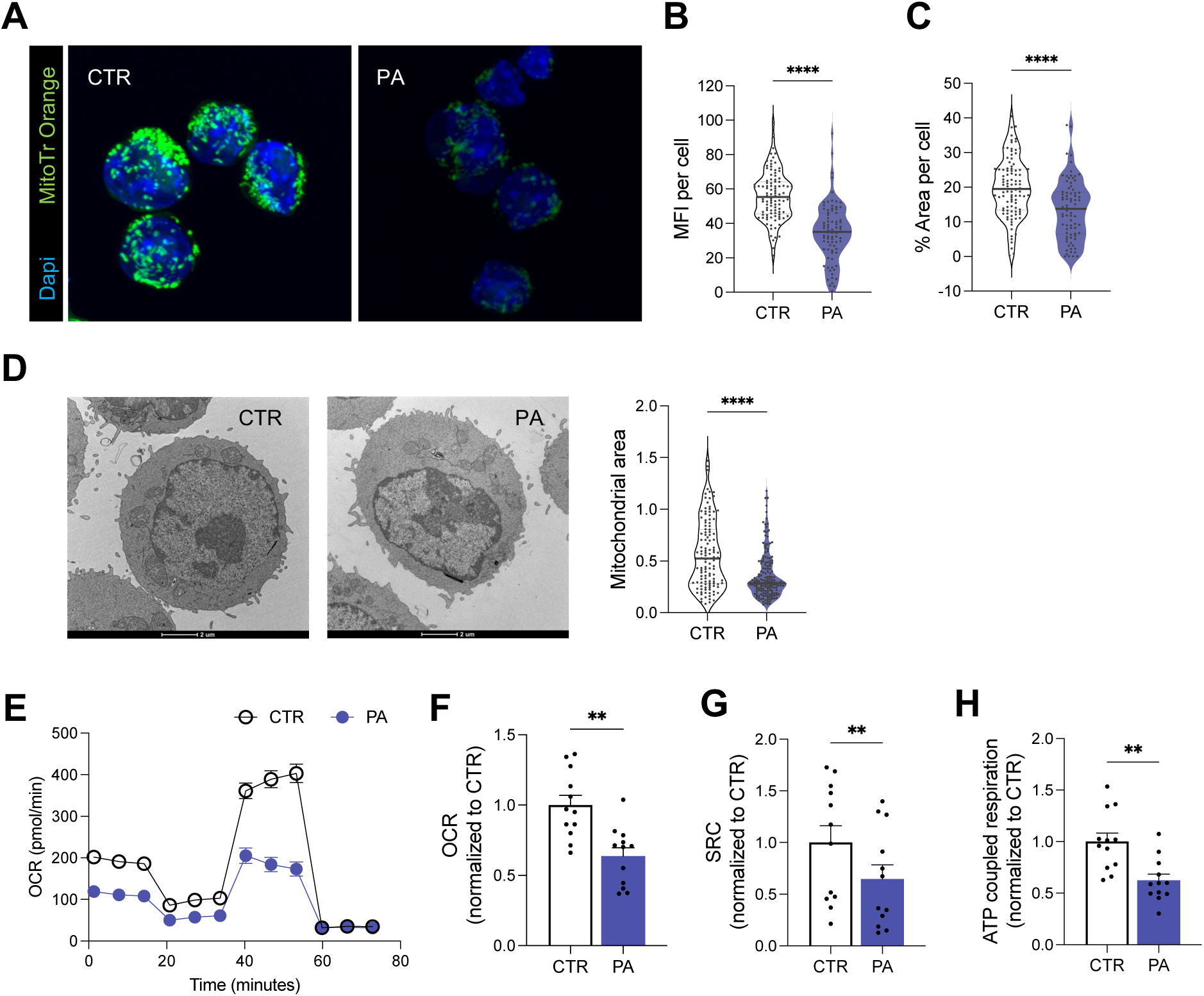
PA disrupts mitochondrial function in CTL. **(A-C).** Mitochondrial profile assessed by multicolor immunofluorescence staining. Representative confocal images with MitoTracker Orange CMTMRos in green, DAPI in Blue and Live staining in purple **(A).** Mean fluorescence intensity (MFI) of MitoTracker orange staining (**B**) and mitochondrial area per cell (**C**) (N=4). **(D).** Representative electron microscopy image and relative mitochondrial area quantification. (N=3) **(E-I).** Bioenergetic profile of CTL±PA during time (**E**) and relative quantification of Oxygen Consumption Rate (OCR) (**F**), Spare Respiratory Capacity (SRC) (**G**) and ATP production (**H**) (N=8).

### Palmitate decreased histone acetylation in CTL

Next, we investigated the mechanism by which PA impairs anti-tumor immune response. PA is metabolized through mitochondrial β-oxidation and contributes to the total pool of cellular acetyl-CoA (AcCoA), whose levels change considerably in response to a series of physiological or pathological conditions (*24*). Thus, we hypothesized that bioenergetic stress and compromised mitochondrial function induced by PA treatment in CTL would affect their availability of AcCoA. Consistent with this idea, AcCoA levels were significantly restricted in PA-CTL (Figure S3A), supporting that the mitochondrial failure induced by PA in CTL leads to a deficiency in the production of AcCoA in CTL.

AcCoA abundance is inextricably linked to histone acetylation and, thus, to cellular activation status (*25*, *26*). Since lipid-derived AcCoA is a major source of cellular histone acetylation (*27*), we reasoned that PA would negatively impact on histone acetylation in CTL by decreasing AcCoA availability. To test this hypothesis, we assess the effects of PA on the epigenetic landscape of CTL. First, by adopting a quantitative mass spectrometry-based proteomic approach (*28*, *29*), we quantified the overall impact of PA on histone post-translational modifications (hPTMs), a fundamental regulatory mechanism of gene expression, which is tightly controlled by enzymes that respond to the availability of metabolic precursors (*30*). In agreement with our hypothesis, PA treatment was accompanied by a significant global decrease of hyperacetylated peptides (i.e. H3K9acK14ac, H3K18acK23ac, and the tri- and tetra-acetylated form of the histone H4 tail), which are typically associated with a transcriptionally active state of chromatin (Figure 3A). Next, the relative abundance of H3K27ac and H3K9ac, which are well-recognized markers of active transcription and T cell activation (*31*, *32*) was striking reduced in PA-CTL when compared to CTR-CTL (Figure 3B-C), which further corroborated the detrimental impact of PA on CTL histone acetylation profile. Removing PA from the culture medium was not sufficient to restore the acetylation levels in PA-CTL (Figure 3D). This observation further implies that the ‘unfitness’ induced by PA is intricately linked to epigenetic modifications, with acetylation emerging as one of the most significantly affected processes. Consistent with this idea, we observed that PA-CTL exposed to trichostatin A (TSA) - a pan-HDAC inhibitor that augment global histone acetylation - reacquire their functional properties, as assessed by restored ability to produce the majority of tested effector cytokines (Figure S3D). Thus, we can conclude that PA-altered metabolism in CTL affects their epigenetic landscape by reducing the levels of chromatin acetylation.

**Figure 3.**
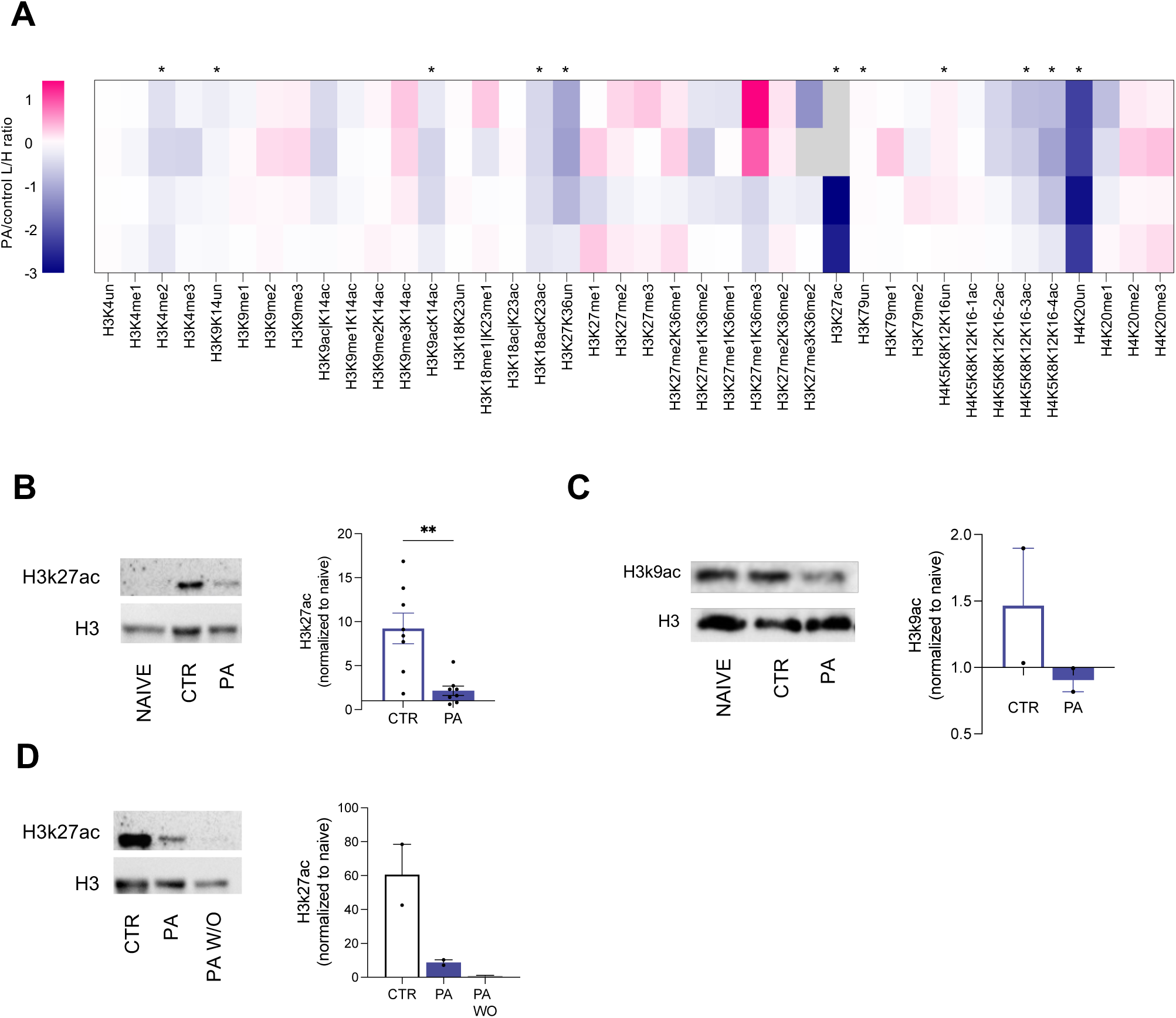
PA reduced histone acetylation in CTL. **(A).** Heatmap representation of histone post-translational modification levels measured by mass spectrometry in CTL±PA normalized on CTR (N=4). The grey color indicates peptides that were not quantifiable. **(B-C).** Western blot analysis and quantification of H3k27ac **(B**, N=7) and H3k9ac **(C,** N=2) in CTL±PA after 48h. **(D).** Western blot analysis and quantification of H3k27ac after 48h of CTL-PA culture followed by 48h with PA (PA) or wash out of PA (PA WO) compared with untreated cells (CTR). (N=2).

### Palmitate impacts chromatin accessibility and gene expression in CTL

Because of the irreversible nature of the effects of PA on CTL proliferation, cytokine production and impaired cytotoxic capabilities, we hypothesized a possible role for PA as an epigenetic regulator of CTL functionality.

Metabolic and epigenetic processes work together to ensure the appropriate CTL cellular responses (*33*) and decreased levels of histone acetylation have been correlated with chromatin inaccessibility and transcriptional repression (*34*, *35*). To assess the contribution of the mitochondrial defects induced by PA on the epigenetic regulation of CTL, we evaluated whether PA treatment was associated with changes in gene expression and/or chromatin accessibility by performing RNAseq combined with assay for transposase-accessible chromatin sequencing (ATACseq) to identify genomic determinants associated with the observed mitochondrial disfunction induced by PA treatment. Gene expression analysis revealed that PA induced significant transcription rewiring, as observed by the 1317 differentially expressed genes (DEGs), of which 468 downregulated and 849 upregulated (log-fold change 0.5, FDR < 0.005; Figure 4A-B and S4A). Gene Ontology (GO) analysis at the functional level revealed that -at functional level-PA treatment induced a significant regulation of genes involved in regulation of apoptosis, mitochondrial activity, lipid catabolism cell cycle and T cell activation/differentiation (Figure 4A-B and S4B-D). This analysis not only confirms the mitochondrial dysfunction phenotype and proliferative defect observed in PA-treated CTLs (Figure 1-2), but also reveals a possible mechanism by which it is perpetrated at the transcriptional level through the effects of PA on histone acetylation. In fact, master regulators of cell cycle (i.e. *Ccne1*, *Mns1*, *Suv39h2*, *Ttk*, *Rbbp8*, *Xlr5b*, *Cntd1)* were significantly downregulated (Figure S4D) upon PA exposure. On the other hand, the central involvement of mitochondria was highlighted by the downregulation of several genes regulating mitochondrial functionality and integrity (i.e. *Nol3, Mpv17l, Pdcd5, Bag3, Timm29, Pnpt1*; Figure S4D). Notably, PA induced higher levels of genes from the mitophagy signature (i.e. *Pink1*, *Adcy1*, *Ambra*, *Adcy3*, *Irgm*, *Lrb*, *Ogt*, *Vps13d*; Figure 4D), luckily as a mitochondria stress cellular response (*36*). Moreover, the induction of carnitine palmitoyl transferase 1 expression (*Cpt1a*), which is the rate-limiting enzyme of mitochondrial b-oxidation, and of very long-chain acyl-CoA dehydrogenase (VLCAD), which catalyzes the first step of the mitochondrial fatty acid oxidation of VL-FAs, indicated the successful delivery of PA and its intracellular metabolism (*37*) (Figure S4D). When looking at GOs enrichment in PA-treated CTL, we observed a deep impact on T cell activity. To corroborate that PA exposure induced a dysfunctional phenotype in CTL, we manually inspected our transcriptomics analysis. Consistent with the repressive role of PA in CTL functional performances, we reported that PA treatment significantly upregulated the expression of genes driving the dysfunctional phenotype in CTL (i.e. *Tox*, *Vsir*, *Tspan32*, *Myb*, *Runx1*, *Lax1*, *Dock2*), while inducing the downregulation of genes involved in critical T cell effector functions (i.e *Ptcra*, *Perf*, *Ifng* and *Gzmb*) (Figure S4D). Our transcriptomic analysis confirms that PA commits CTL to dysfunction by perturbing CTL mitochondrial functions and transcriptional profile; thus confirming that PA acts as a negative regulator of CTL biology.

**Figure 4.**
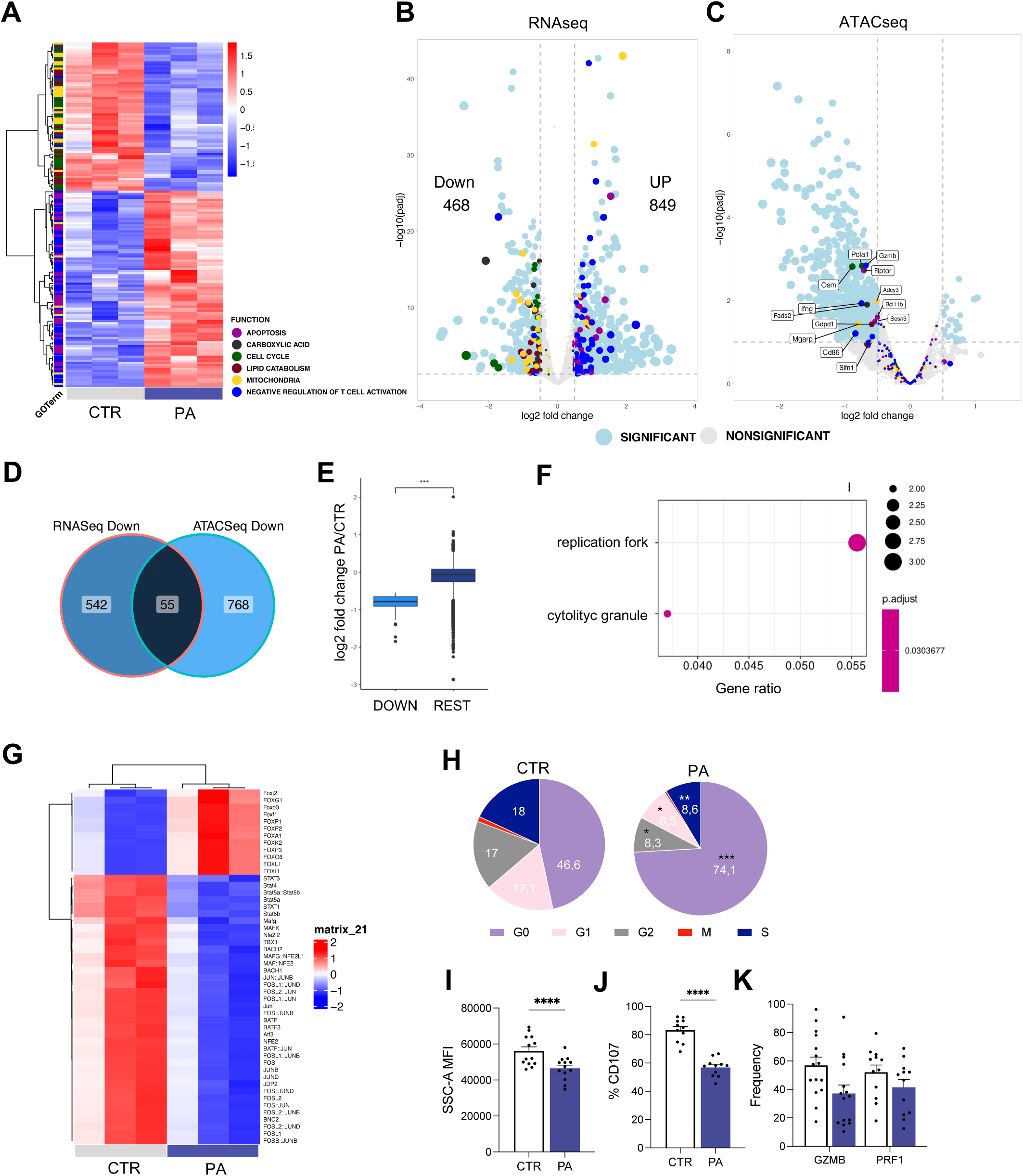
PA decreases chromatin accessibility in CTL. (**A**) Heatmap representing genes from the top enriched GO Terms color-coded as shown in the legend. Rlog-transformed values were normalized row-wise (z-score legend) and each gene was colored by its respective GO Term. (**B**) Volcano plot showing differentially expressed genes (DEGs) in CTL (negative logFC) vs PA (positive logFC). Dots are colored by GO terms and their sizes represent the log-fold quantity. (**C**) Volcano plot of ATACSeq promoters labeled by GO Terms shown in A. Genes downregulated in both RNASeq and ATACSeq are labeled. (**D**) Venn Diagram showing the promoters of common genes downregulated by PA in RNASeq and showing inactive chromatin in ATACSeq. (**E**) Boxplot showing the LogFold values of the promoters of the genes downregulated in RNASeq and ATACSeq vs all the others in ATACSeq dataset. (**F**) GO terms enriched on significantly dowregulated gene promoters in PA-CTL. (**G**) Top 50 enriched transcription factors (TF). (**H**) Pie chart of cell cycle phases in CTL±PA (N=4) (**H**) Representative histogram and quantification of SSC-A (N=13). (**I**) Quantification of CD107 expression (N=11). (**J**) Quantification of Cytokines production: GZMb and PRF1 (N=12).

Dysfunctional CTL are defined by specific chromatin states and epigenetic program (*38–43*). To elucidate the epigenetic states associated with PA-mediated CTL dysfunction, we employed chromatin accessibility with the assay for transposase-accessible chromatin sequencing (ATAC–seq (Figure 4C and S4F-H). Overall, PA treatment induced an important chromatin remodeling, with the majority of the significant changes being towards a more compact chromatin state. In fact, only 72 genomic regions were characterized by increased chromatin accessibility following PA-treatment, whereas 5941 regions significantly became less accessible (Figure 4C).

This result is in line with the fact that PA induces loss of acetylation, which is often associated with open chromatin states. In agreement with that, PA-treated chromatin showed a tendency of overall more compactness with tetra nucleosome being much more abundant than in the control condition (Figure S4I). GO analysis of genes showing a closer chromatin state revealed they were mainly involved in mitochondrial, cell-cycle and T cell activation (Figure 4C), supporting our hypothesis of a specific PA-driven T-cell dysfunction remodeling at the chromatin level. Next, we sought to investigate whether a common core could be identified between the ATACseq and the RNAseq analysis, which would probably represent genes whose control is deeply affected by PA treatment. To this aim, we intersected expression and epigenetic changes for the top scored downregulated genes at the ATAC (Promoter region) and RNAseq levels. 56 genes appeared to be regulated at both levels (Figure 4D and S4I). Strikingly, cellular compartment (CC) GO analysis highlighted an enrichment of terms specifically related to replication fork formation (*Rfc1*, *Pola1* and *Rfc4*) and cytolytic granules (*Gzmb* and *Prf1*) ((Figure 4E-F), suggesting that the chromatin remodeling operated by PA treatment affected deeply DNA replication and the cytotoxic effector capabilities of CTL. This was functionally investigated by analyzing cell cycle profile and granulitic abilities of CTL activated in the presence or absence of PA. Upon proper T cell stimulation, 18% of CTL were in S phase, indicating that they were actively synthesizing DNA and undergoing cell division. However, upon PA exposure, only 8,6% of the cells progressed into S phase and most of the PA-CTL remained in G0 phase (Figure 4H), indicated that the treatment with PA arrested CTL in the G0/G1 phase of the cell cycle, in line with their DNA duplication defect observed when intersecting ATAC and RNAseq. Likewise, PA-CTL, compared to CTR-CTL, displayed a lower granularity (SSC, Figure 4I) twinned with an impaired ability to degranulate upon antigen-restimulation, as indicated by the lower amount of the granular membrane protein CD107 (Betts MR et al. 2003) (Figure 4J) and decreased ability to secrete granzyme-B and perforin (Figure 4K). These defects provided an explanation of the impaired anti-tumor potential observed in PA-CTL (Figure 1E). Thus, we can conclude that exposure of CTL to PA that mediates the CTL dysfunction by restraining chromatin accessibility at genes crucial to sustain CTL cell cycle progression and granulitic functions, both essential to coordinate the anti-tumor immune response.

We next analyzed transcription factor (TF) binding motifs represented in chromatin regions that became significantly less accessible in CTL exposed to PA (Figure 4G). Strikingly identified transcription factors that were all members of the transcription factor activator protein-1 (AP-1), a master regulator governing crucial aspects of CTL biology such as activation, differentiation, anergy and exhaustion (*44*, *45*). In particular, we identified components of the Jun (c-Jun, JunB, JunD), Fos (c-Fos, FosB) and the activating transcription factors, ATF (LRF1/ATF3, BATF, JDP2) protein families, which have been shown to function downstream the TCR and induce expression of IL2 and other effector molecules in activated CTL, as well as promote chromatin accessibility at the enhancers of genes coding for effector functions in CTL. These data indicate that PA may also restrict CTL activity by impeding proper CTL activation and, therefore, limiting the function of several members of the AP1 complex, required to keep an open chromatin state during T cell activation (*46*). In summary, our findings reveal that PA induces a state of metabolic dysfunction in CTLs, characterized by profound epigenetic disturbances which impede the appropriate activation of CTL, thereby uncovering a previously unrecognized role of PA. This role establishes a critical mechanistic link among metabolic pathways, epigenetic regulation, and the functional differentiation of CTL, offering new insights into the intricate interplay between cellular metabolism and immune response.

### SPHK2 regulates functionality and metabolic commitment of PA-CTL

Next, to gain mechanistic insights on PA-induced CTL “unfitness”, we aimed to identify metabolic features associated with PA exposure. To this end, we performed an unbiased metabolomic profiling of PA- and CTR-CTL. When compared to CTR-CTL, PA-CTL showed an enrichment in a subset of palmitate-containing lipids (Figure 5A), suggesting that PA is up-taken and metabolized by CTL, in accordance with CPT1a and VLCAD upregulation (Figure S4D). In particular, PA increased levels of sphinganine, sphingosine and sphingomyelin in CTL (Figure 5A and S5A), suggesting that PA induces a positive modulation of the sphingolipids (SL) metabolic pathway in CTL. Accordingly, a deeper analysis of the lipidomic profile of PA-CTL recorded broad changes in the abundance of different types of SL, including ceramides (Cer), hexosylceramides (HexCer) and phosphoethanolamine (PE), all important intermediate in the SL metabolic cascade (Figure 5B). Of note, the lipidomic analysis of PA-CTL upon PA exposure recorded broad changes in the abundance of several phospholipids (PLs), such as lysophosphatidylcholines (LPCs), phosphatidylcholine (PCs), phosphatidylethanolamines (PEs) and phosphatidylinositols (PI) (Figure 5B). Such changes in PC composition are predicted to alter the maintenance of membrane integrity. Furthermore, multiple lipid classes enriched in PA-CTL displayed an overall increase in the degree of saturation of the acyl-chains (Figure S5B), which has been predicted to increase membrane rigidity (*6*, *47*). Accordingly, when we probed the functional consequences of PA-induced modification in the lipidome of CTL by calculating their respective membrane fluidity, we reported that membranes of PA-CTL were less fluid in comparison to those of CTR-CTL (Figure S5C), a difference that provides an explanation for the reduced activation and function observed in PA-CTL. Thus, PA guides a variation of the lipid composition of CTL cellular membranes by increasing their rigidity and promoted the formation of sphingolipid mediators, which might be responsible for mediating the PA-mediated dysfunctional program in CTL. Sphingolipids (SL) and ceramides have been both reported to cause severe defects in cellular growth, differentiation and proliferation in nonimmune cells (*48–50*); thus, we questioned whether the mechanism behind PA-induced CTL dysfunction might be routed in SL elevated levels in PA-CTL. Based on earlier work (*51*) and on the metabolic profile associated with PA, we reasoned that the PA-induced mitochondrial dysfunction in CTL could be associated with the ability of PA to promote the activity of sphingosine kinase 2 (SPHK2), a key enzyme in the sphingolipid metabolic pathway that catalyze the formation of sphingosine 1-phosphate (S1P) (*52*, *53*) Hu et al J of Lipid Research 2009) but also convert S1P back to sphingosine and then to ceramide, thus further promoting mitochondrial damage (*54*) (Figure 5C). In accordance, the levels of S1P were significantly higher in PA-CTL compared with CTR-CTL (Figure 5D). Thus, the elevation of PA levels in CTL led to an increase in the levels of S1P, a lipid molecule with bioactive properties regulating a variety of cellular processes such as cell death, movement, proliferation, differentiation (*55*, *56*) and a major regulator of T cell biology and function (*57–60*).

**Figure 5.**
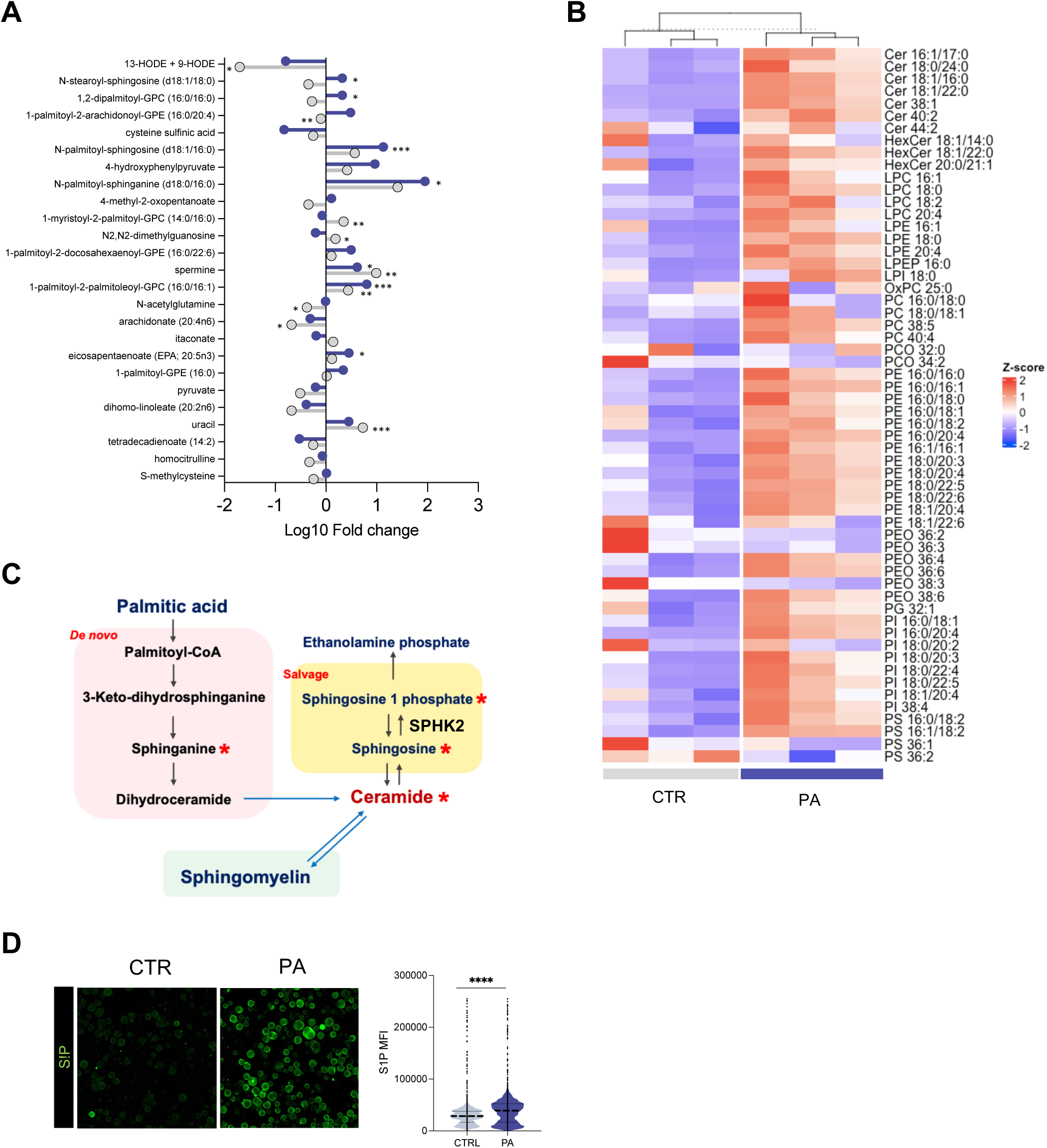
PA-induced dysfunction is with SPHK2 upregulation in CTL. **(A).** Lollipop representation of significantly regulated metabolic compounds in CTL±PA in respect to naïve CTL (N=8). **(B).** Heatmap from LS-MS lipidomic analysis (N=3). **(C).** Graphical model of sphingosine metabolism. Stars indicate upregulated compounds in PA-CTL. **(D).** Immunofluorescence imagines and relative quantification of S1P in CTL±PA. S1P in green (N=4).

Moreover, treatment of PA-CTL with bezafibrate to lower the levels of generated very-long chain FAs and S1P production, as well as fingolimod to suppress the action of S1P (*61*), did not improve the mitochondrial defects associated with PA treatment (Figure S5D); further supporting a possible implication for SPHK2 in mediating PA-induced CTL metabolic and functional “unfitness” by increasing SL and ceramide accumulation.

### *Ex vivo* inhibition of SPHK2 restores functional fitness in PA-CTL

SPHK2 plays an important role in mitochondrial dysfunction and its targeting is a promising strategy to enhance mitochondrial function and metabolic fitness (*62*, *63*). Having established a link between PA exposure, SPHK2 activity and mitochondrial dysfunction, we next aimed to test whether pharmacological SPHK2 inhibition (SPHK2*i*) might counteract the CTL dysfunction induced by PA, using a novel, highly specific SPHK2 inhibitor, ABC294640 (*64*), emerging as a promising option for treating different type of cancers (*65*).

SPHK2 has been shown to negatively regulate CD4 T cell function during viral infection (*57*); however, its role on shaping activity and differentiation of CTL remains to be investigated. Thus, we first evaluated the effects of SPHK2*i* on CTL performance. After stimulation, CTL got fully activated CTL in presence of SPHK2*i*, as assessed by cell surface expression of activation markers (i.e CD44, Figure S6A) - without affecting their viability (Figure S6B) - and we did not observed differences neither in the degree of proliferation (Figure S6C) nor in the frequencies of effector/memory population (Figure S6D) between CTL and SPHK2*i*-CTL. Moreover, upon TCR restimulation, effector cytokines - IFNγ and TNFα (Figure S6E) and cytotoxic molecules – GZMB and PRF1 (Figure S6F) – were similar between CTR and SPHK2*i*-CTL, resulting in an equal ability to kill a cognate tumor-cell line (Figure S6G). In all, these results indicate that SPHK2i did not alter activation, differentiation and effector functions of CTL.

Subsequently, we sought to examine whether targeting SPHK2i may counteract the PA-induced metabolic dysfunction and, thus, ameliorate CTL anti-tumor potential, especially in the context of lipid-rich TME. To this end, we first assessed how SPHK2*i* affects PA-induced dysfunction in CTL. SPHK2*i* proved able to restore the PA-induced mitochondrial defects as demonstrated by the recovery in both mitochondrial function and mass (Figure 6A-C), which is mirrored also by the reacquired ability to engage mitochondrial OXPHOS (Figure 6D) as well as enhanced ATP generation (Figure 6E), reaching the levels displayed by the optimally activated CTR-CTL. Thus, SPHK2*i* in PA-CTL leads to the recovery of an optimal bioenergetic profile, reverting the metabolic “unfitness” induced by PA in CTL.

**Figure 6.**
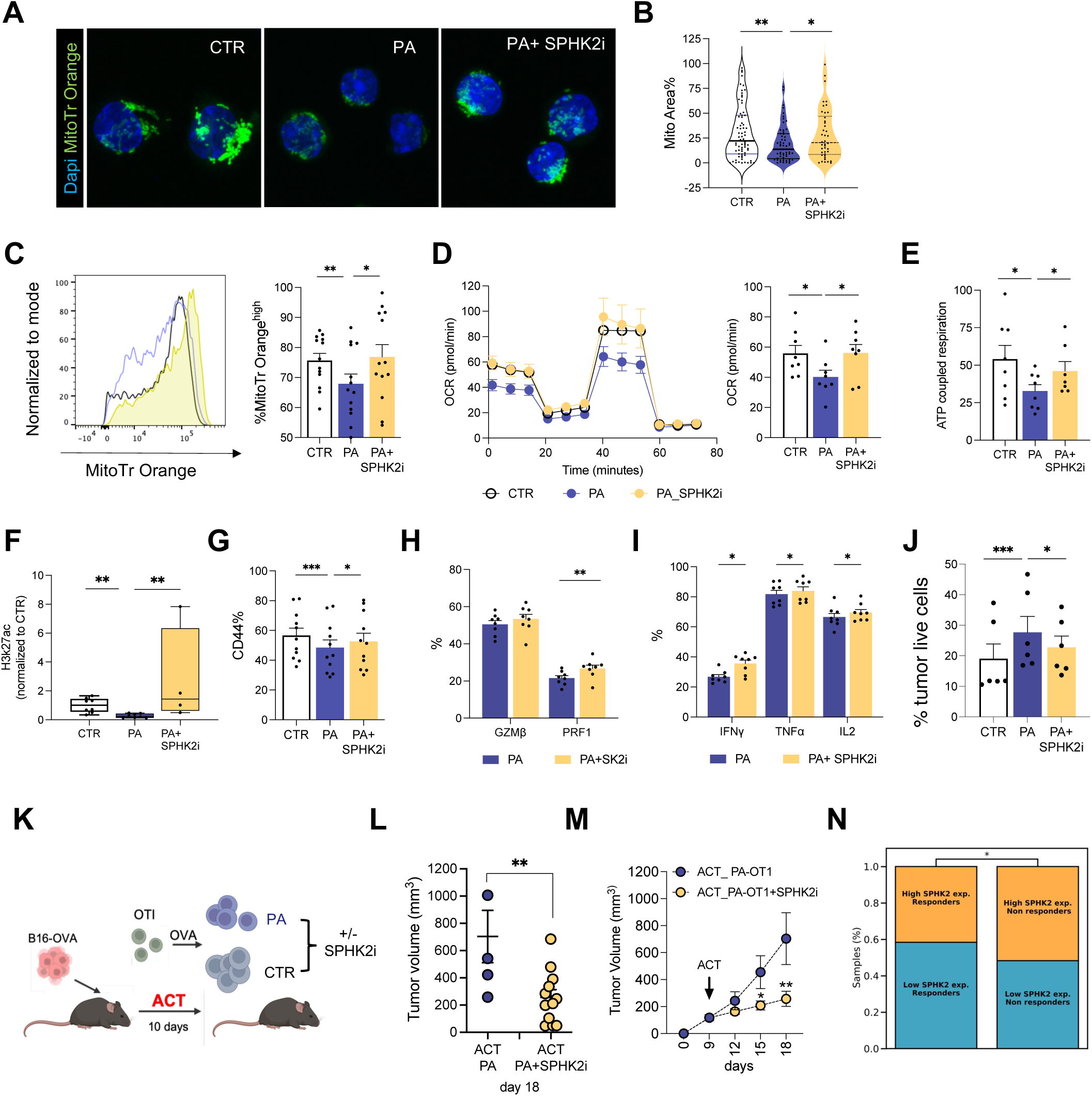
SPHK2 inhibition reverts PA-induced dysfunction. **(A-C).** Mitochondrial potential in CTL±PA alone or in combination with SPHK2i assessed by multicolor immunofluorescence (MitoTracker Orange CMTMRos in green, DAPI in Blue and Live staining in purple, **A-B**) or flowcytometry (N=11) (**C**). **(D-E).** Bioenergetic profile and quantification of OCR (**D**) and ATP production (**E**) (N=8). **(F).** Relative abundance of H3k27ac assessed by WB (N=4) **(G-J).** Quantification of CD44 expression (**G**, N=9), cytokines production (**H-I**, N=9) and CTL killing activity against B16OVA (**J,** N=4). **(K).** Experimental design of in vivo adoptive cell therapy with PA-CTL ±SPHK2i. **(L).** Growth curves of B16OVA tumor treated with PA-OTI ±SPHK2i (N=4/15). **(M).** Tumor volume of B16OVA tumor bearing mice treated with PA-OTI ±SPHK2i (N=4/15). **(N).** Association between SPHK2 expression and response to ICB therapy on five aggregated datasets from lung, skin and kidney cancer from the Tumor Immunotherapy Gene Expression Resource database (Chi-square, p = 0.04).

Next, we tested whether -by reestablishing the mitochondrial functionality in PA-CTL-SPHK2*i* was capable to abolish the acetylation defects correlated with PA-induced mitochondrial dysfunction, that we showed be responsible for the activation and differentiation defect in CTL. Strikingly, H3K27 acetylation levels were significantly increased in PA-CTL upon SPHK2*i* (Figure 6F and S6H), providing evidence that SPHK2i might counteract the CTL metabolic dysfunctional state by correcting the epigenetic acetylation defects induced by PA treatment. Thus, we next hypothesized that the recovered mitochondrial functionality in concert with restored acetylation levels after SPHK2*i* treatment would restore CTL functionality under PA conditions. Accordingly, flow cytometry profiling revealed that SPHK2*i* significantly enhanced the ability of PA-CTL to get activated (Figure 6G) produce effector cytokines (Figure 6H) as well as cytotoxic molecules (Figures 6I). Most importantly, this results in an ameliorated anti-tumor potential - as indicated by the improved tumor killing ability in PA-SPHK2*i* CTL (Figure 6J). In all, these data suggest that SPHK2i operates a concert metabolic and epigenetic rewiring which re-energize CTL and improve their effector functions. In this context, to strengthen the potential role of SPHK2 inhibition as a strategy to re-establish CTL functional fitness, we isolated PDA infiltrating CTL from pancreas of KC mice of 20 weeks and treated them *ex vivo* with or without SPHK2i. Based on our previous study (*12*), we reasoned that at the time of Late PanIN lesions, infiltrating CTL would have experienced prolonged lipid exposure and defective phenotype analogous to T cells cultured in vitro with PA. Treating PDAC-infiltrating CTL with SPHK2i *ex vivo* significantly increased their mitochondrial function (Figure S6I), proving that counteracting SPHK2 activity promotes metabolic responsiveness in CTL isolated directly from the lipid-rich TME.

To finally prove that the recovered fitness induced *in vitro* in PA-CTL by SPHK2i enable them to better adapt to the inhibitory TME and mediate a durable tumor control in the *in vivo* setting, we employed an ACT model for established B16-OVA melanoma tumors. We generated OVA-specific T cells *ex vivo* in the presence or absence of PA and treated or not with SPHK2i and injected them intravenously in tumor-bearing mice (experimental scheme in Figure 6K). SPHK2i enabled PA-CTL to control tumors as efficiently as CTR-CTL (Figure 6L and S6J), resulting in a significantly decreased tumor volume compared to tumor-bearing mice adoptively transferred with PA-CTL (Figure 6M), which, in accordance with their dysfunctional metabolic and functional status, were associated with a more rapid and worse tumor progression (Figure 6L). Thus, we underscore a new role for SPHK2i in CTL anti-tumor response being able to counteract metabolic and functional defects in CTL towards a functional phenotype able to operate tumor debulking. Finally, we explored whether SPHK2 expression correlates with reduced survival prognosis in human cancers by analyzing the survival of high vs low SPHK2 expression samples in 29 TCGA projects. SPHK2 expression was associated with lower survival probability in melanoma patients and across multiple cancer types (Figure S6K). No cancer type shows an association of high SPHK2 expression with better survival. Having proved SPHK2 as a key regulator of anti-tumor potential of CTL in metabolic stressed conditions - such as lipid-rich exposure – we hypothesized that SPHK2 expression might reduce CTL functions in human tumors. Thus, we evaluated whether SPHK2 expression was associated with therapeutic outcomes in cancer patients undergoing immune checkpoint blockade (ICB) therapy. We retrieved five datasets from lung, skin and kidney cancer treated with anti-PD1 drugs from the Tumor Immunotherapy Gene Expression Resource database (TIGER) (*67*). When testing for association between low SPHK2 expression and PD1 response, we saw that in every single dataset, there was a higher fraction of lowly SPHK2 expressing samples among responders as compared to non-responders. Due to small study sizes, we pooled studies by stratifying first patients within each study be lower vs higher than median SPHK2 expression and then tested again for association with response in the aggregated cohorts. We observe a statically significant association between low SPHK2 expression and response to ICB (p = 0.04; Chi squared test, Figure 6N). Overall, these findings identify SPHK2 inhibition as a new strategy to energize CTL, possibly improving CTL anti-tumoral responses, with a particular relevance for tumors characterized by a lipid-rich TME. In all, our study identified PA as a unique metabolite at the intersection of metabolism and epigenetic regulation of CTL differentiation. PA prompts a mitochondrial dysfunctional state via SPHK2 regulation that demotes histone acetylation and chromatin accessibility in CTL, impairing their proper activation and differentiation. Consistently, pharmacological inhibition of SPHK2 restored CTL mitochondrial fitness, effector functions and anti-tumor potential (Figure 7).

**Figure 7.**
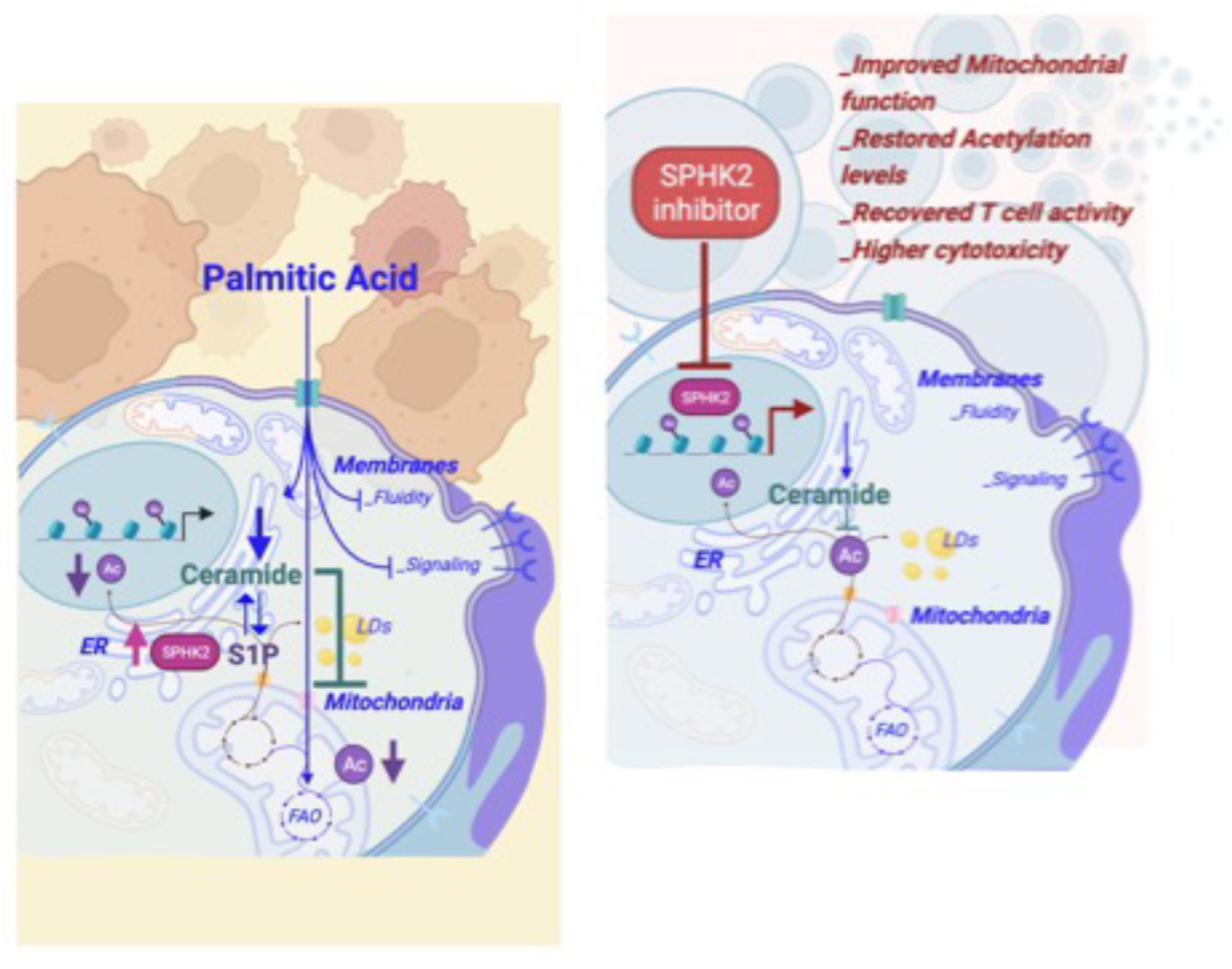
Graphical model.

## DISCUSSION

In this study, we discovered an unknown dimension of PA’s role in cancer pathogenesis. Beyond its established pro-tumoral mechanism (*17*, *18*), we identified a critical, pro-tumoral extrinsic function of PA in significantly undermining CTL-mediated anti-tumor immunity. Here, we propose that PA can foster tumor growth not only through tumor cell intrinsic mechanisms but also by directly preventing CTL anti-tumor potential. This dual functionality of PA not only expands our understanding of the tumor progression process but also opens new, groundbreaking avenues for therapeutic intervention, potentially transforming the landscape of cancer treatment.

Lipids, as major nutrient source in TME, are exploited by cancer cells to support their growth and dissemination (*6*). In such lipid-rich TME, tumor-infiltrating CTL can potentially use lipids as an alternative source of energy to fuel their effector functions in glucose-deprived TME. However, altered lipid levels leads to CTL dysfunction (*11*, *12*, *15*, *68*). Analysis of the metabolic composition of different solid tumors (*12*, *17*) has revealed PA-enriched TME and metabolic dysfunctional signatures in tumor-infiltrating CTL. Mechanistically, we unveil here that PA alters CTL metabolism by disrupting mitochondrial functionality. Decreased mitochondrial activity caused by PA had previously been reported in other cellular contexts, but our work shows that PA can additionally have a novel, direct and immediate effect on mitochondrial metabolism that prevents CTL activation. Future investigation is required to determine whether PA-induced metabolic unfitness expands beyond CTL and impact on other immune cells within the TME.

Lipid metabolism plays a central role in regulating CTL activation and differentiation. As effector, CTL upregulated lipid anabolism to meet their growing biomass requirements, while memory CTL burn fats to stay in a relatively quiescent and persistent state (*10*, *69*). Furthermore, CTL enhance lipid catabolism to preserve their anti-tumoral functions (*70*). Our findings showed that PA diverts CTL cellular metabolism towards sphingolipid metabolism, resulting in mitochondrial damage and cell-cycle arrest - as previously shown in other cellular contexts (*50*, *54*). Additionally, we identified an unknown role for lipid metabolism in regulating the process of cell-cycle progression, proliferation and degranulation processes in CTL. Thus, rather than a dysfunctional differentiation program, our data points at a T cell activation defect in PA-rich conditions, expanding upon previous work and demonstrating the role of lipid metabolism in regulating T cell effector functions and differentiation (*9*, *71*).

The deregulation of lipid metabolism is often accompanied by the transcriptional reprogramming of dysfunctional CTL, proving that metabolic alterations are transcriptionally regulated (*12*). In addition, lipids can directly interact with epigenetic modifying enzymes or their substrates to regulate CTL differentiation (*72*, *73*). Adding to this, we functionally link the metabolic perturbations induced by PA with specific epigenetic alterations that, ultimately, lead to functional defects in CTL. Thus, we identified a previously unrecognized role for lipids in directly shaping CTL fate and function through epigenetic modifications. As lipids hold a cell-type-specific modulation of mitochondrial metabolism and, as a consequence, cell functionality (*9*), it will be important in future investigation to dissect the context-dependent role of lipids in epigenetic regulation. Hence, decoding the epigenetic machinery of CTL infiltrating lipid-rich TME might offer an insight as to how specific lipids support cancer cells while impairing immune cells functions.

Metabolic reprogramming is a unique strategy to promote survival and functionality of CTL in the metabolically hostile TME (*74*). Here, we identified SPHK2 as a metabolic modifier that regulates mitochondrial fitness in CTL. *In vitro* inhibition of SPHK2 protects CTL from the lipid-induced dysfunction by promoting a potent mitochondrial rewiring in CTL, which is able to restore functional and metabolic fitness in lipid-exposed dysfunctional CTL. More work will need to be done to determine whether SPHK2i in CTL acts by inhibiting the kinase activity of SPHK2 or by affecting intra- and extra-cellular levels of S1P as well as how its inhibition impact on CTL lipid composition in relation to their mitochondrial rewiring.

Obesity and diets enriched in fat (HFD) (*75–77*), especially saturated ones, potentiate the pro-tumoral effects of lipids, limiting nutrient availability for CTL, thus impairing their function (*78*). With PA being the primary saturated fatty acid in most diets, our results highlighted an unprecedented need for dietary interventions to modulate feeding regime with the perspective to prevent tumor spreading while favoring anti-tumor responses and, thus, improve the quality of life of cancer patients.

## MATHERIAL AND METHODS

### Animal studies

All animal experiments were conducted in accordance with European Union Guideline on Animal Experiments under the protocol number #660/21. The animals were kept in a pathogen-free facility, following a 12-hour light/dark cycle, and were provided with unlimited access to standard rodent chow and water. The KC (p48Cre;LSL-KrasG12D) mouse, generated as previously described (*79*, *80*), were housed at European Institute of oncology (Milan, Italy). KC animals were euthanized at 10-20 weeks (Early lesion) and 20-30 weeks (Late lesion). For other mouse experiments, we used 8-10 weeks old C57BL/6J and OT1 female mice. The number of animals has been indicated in the respective figure legends.

### Isolation of T cells from tissues and activation

Mouse T cells were harvested from pancreas, spleen, and lymph nodes (inguinal and axillary) harvested from mice under sterile conditions. Briefly, the spleens and lymph nodes were minced in RPMI, 10% FBS, 1% Pen/ Strep, and the resulting cells were filtered through a 40-µm filter. To eliminate red blood cells, ACK lysing Buffer (Gibco) was used. For tumor-infiltrating lymphocytes, pancreas were collected, cut into small fragments, and exposed to a mixture of 0.8 mg/ml collagenase IV and trypsin inhibitor from Sigma-Alrich in 1× HBSS (Thermo Fisher Scientific) for 20 minutes at 37°C. The tumor fragments were then homogenized, washed, and strained through a 70-µm filter. CD8+ T cells were sorted by negative selection with CD8a+ T Cell Isolation kit Mouse (Miltenyi) following the manufacturer’s instructions or by FACS sorting. CD8+ T cells were resuspended at 1×106 cells/mL in complete T Cell Media (TCM, RPMI, 10% FBS, 1% NaPy, 1% Glutamine, 1% Pen/ Strep, 0,1% b-mercaptoethanol) supplemented with 100U/mL IL2. For C57J/B6 mice derived cells polyclonal activation was achieved using 5 μg/mL plated-bound aCD3 and 0,5 μg/mL soluble aCD28, while for OT1 mice, media was complemented with 2 μg/mL of SIINFELK peptide. For the characterization of the effect of palmitate on CD8+ T cells functionality, media was further supplemented with Palmitate-BSA conjugated (PA) at a concentration of 100 μM. For the inhibition of SPHK2, T cell media was supplemented with ABC294640 at a concentration of 12,5 nM. Unless otherwise specified, cells were cultured 48 hours at 37°C, 5% CO2 and 90-95% humidity for a pH neutral environment.

### Tumor processing

Tumors were excised from mice at the indicated time point post-implant. After harvesting, tumors were chopped and digested in DMEM, 1 mg/ml collagenase A, 100 µg/ml DNase I for 30 minutes at 37°C. Tumor digestion was strained through a 100- and 70-µm filter. Cells were washed and counted. Cells (1×10^6^) were further stained for flow cytometry analysis.

### Immunophenotype and flow cytometry

Flow cytometry was employed to characterize Palmitate-treated CD8+ T cells, following standard protocols. In brief, 1×10^5^ - 1×10^6^ cells were collected at specified post-activation time points, washed with PBS, and then suspended in FACS buffer (PBS, 2% FBS, EDTA 2mM). The cells were subsequently stained for surface markers at 4°C for 20 minutes. For intracellular staining of cytokine production, the cells were incubated for 4 hours in complete RPMI with a 1:1000 dilution of GlogiPlug (BD), either with or without secondary stimulation using PMA/Ionomycin (at 20 ng/mL and 1 μg/mL, respectively) at 37°C. The cells were fixed and permeabilized using the Fix/Perm kit BD (560409) as per the manufacturer’s instructions. Intracellular staining was then carried out in permeabilization buffer for 2 hours at 4°C. Additionally, CFSE, MitoTRK Orange and Green stainings were conducted following the manufacturer’s guidelines. For SPHK2 staining, cells were first stained for surface markers, then fixed and permeabilized using the the Fix/Perm kit BD following the manufacturer’s instructions. A primary antibody (SPHK2, BS-2653R, Bioss) was used for the first staining in permeabilization buffer for 1 hour, followed by a wash in PBS and a 1-hour incubation with the secondary antibody in permeabilization buffer. The samples were acquired using a BD FACS Canto II or Celesta.

### Immunofluorescence and confocal microscopy

Confocal microscopy with fixed cells was performed on a LEICA SP8 FSU confocal (Leica Microsystems GmbH, Wetzlar, Germany) microscope (Imaging Unit, IEO) with a 40X/1.3, 63X/1.4 or 100X/1.47 oil-immersion objectives. Cells were pre-stained with Mito tracker Orange and a fixable viability dye; cells were allowed to attach by gravity on culture slides coated with poly-d-lysine; and then fixed with 4% paraformaldehyde and permeabilized with 0.1% Triton X-100. Cells were permeabilized for 1hour in blocking buffer (PBS, Triton 0,1%, 3% BSA and 5% FBS) and SPHK2 staining was performed overnight at 4°C in blocking buffer. Cells were washed with PBS and stained with DAPI and the indicated secondary antibody for 2 hours at room temperature. For static killing experiments, B16-OVA cells were plated on a Lab-Tek II chamber slide and left to adhere for 6 hours prior to co-culture with activated CD8+ T cells. OT1 CTL activated in the presence or absence of Palmitate for 48 hours were then counted and added to the chamber in a Effector: Target (E:T) ratio of 1:1. 24 hours post-co-culture, chambers were washed, stained with aCD8 for 30 min at 4°C in PBS; fixed and permeabilized as previously described. Cells were then stained with primary anti-body aCaspase 3 overnight at 4°C, labelled with the appropriate secondary antibody and DAPI for 2 hours at room temperature.

For the colocalization analysis, DAPI, anti-SPHK2 labelled with an AlexaFluor488 labelled secondary antibody, MitoOr. and L/D signals were excited with 405 nm, 488 nm, 568 nm, 633 nm laser lines and the emitted fluorescence acquired in the following acquisition windows: 410-512, 495-545, 566-668, 638-775 nm, respectively. The Z-stacks were deconvolved with Huygens software and the central section of each cell was used to calculate the percentage of colocalization. A macro in Fiji/ImageJ124 has been created. Briefly, region of interests (ROIs) including only one cell were split in the single fluorescent channels, the DAPI, SPHK2 and MitoOr. channels were segmented using the Moments and Otsu algorithm, respectively, and the area occupied by the two signals was calculated. The colocalization area was calculated as the interception between the IP3R and the MitoTracker Orange binary images and expressed as Mean fluorescence intensity MFI respect to the total area occupied by the mitochondria and nuclei respectively.

### Electron microscopy

To prepare the samples for Transmission electron microscopy (TEM) analysis, they were fixed using a solution containing 3% glutaraldehyde and 2% paraformaldehyde in 0.1 M cacodylate buffer with a pH of 7.3 for 1 hour. After fixation, the samples underwent a series of treatments, including washing and treatment with 0.1% Millipore-filtered cacodylate-buffered tannic acid, followed by post fixation with 1% buffered osmium tetroxide for 30 minutes. Subsequently, the samples were stained with 1% Millipore-filtered uranyl acetate. Dehydration was performed using increasing concentrations of ethanol, and the samples were then infiltrated and embedded in LX-112 medium. Finally, polymerization was carried out at 60°C for 2 days. Ultrathin sections were obtained using a Leica EM FC7 ultramicrotome. After staining with uranyl acetate and lead citrate, the sections were analyzed using a Leo 912AB transmission electron microscope (Carl Zeiss). Images were acquired with a 2K bottom-mounted slow-scan Proscan camera controlled by EsivisionPro 3.2 software. For morphometrical analysis, 21 random whole cellular profiles were captured for each experimental condition. These profiles were stitched together from several images acquired at a nominal magnification of 6,300× (pixel size, 1.77 nm). ImageJ was used evaluate the mitochondria size.

### RNA sequencing

The RNA of CD8+ T cells, both treated with and without PA for 48 hours, was extracted from purified cells using RNeasy Mini kits (Qiagen). Quality control was conducted using a a 6000 bioanalyzer (Agilent) and the RNA concentration RNA concentrations were determined using a Qubit 2.0 Fluorometer (Thermo Fisher Scientific).

After overall quality control with FastQC and adaptor removal with TrimGalore (v 0.6.10), paired-end reads, each 51 base pairs in length, were aligned to the GRCm39 mouse genome (release 109). The indexing of the genome was carried out with Salmon, and both the alignment and quantification of read counts were performed with the default parameters. Subsequent analyses were conducted within the R environment (v 4.3). Transcript abundance at the gene level was estimated with the tximport package, followed by Deseq2 to perform differential expression analysis comparing three replicates of Palmitate-treated cells ith their corresponding controls. Genes with very low read counts (minimum 10 reads in 3 replicates) were discarded before differential expression analysis. The criteria for significance in the differential expression analysis were set at a minimum log-fold change of 0.5 and an adjusted p-value less than 0.05, following the Benjamini-Hochberg correction (FDR). Gene annotation was done using the EnsDb.Mmusculus.v79 package and Gene Ontology (GO) term enrichment analysis was carried out with the ClusterProfiler package with default parameters. Redundant terms were eliminated from the enrichment results using the GOSemSim package with simplify function. For data visualization, volcano, radars and dot plots were generated using the ggplot2 package, while heatmaps were generated with complexHeatmap package (*81*) in the R environment.

### ATAC-seq

Libraries were created following a modified protocol from Buenrostro et al (*82*). In summary, 50,000 FACS-purified CD8+ T cells per subset were washed with PBS and suspended in 50 μl of lysis buffer (10 mM Tris-HCl pH 7.4, 10 mM MgCl2, 0.1% IPEGAL CA-630). Nuclei were collected by centrifugation for 10 minutes at 500 g and then resuspended in a final reaction volume of 50 μl, containing 1 μl of Tn5 transposase (produced in-house), 10 μl of 5x transposase buffer (50 mM Tris-HCl pH 8.4, 25 mM MgCl2), and 39 μl of ultrapure water (Milli-Q). The reaction was incubated with mixing at 300 rpm for 30 minutes at 37°C. Subsequently, 10 μl of clean-up buffer (900 mM NaCl, 30 mM EDTA), 5 μl of 20% SDS, 0.7 μl of ultrapure water (Milli-Q), and 4.3 μl of proteinase K (18.6 μg/μl, Thermo Fisher Scientific) were added, and the mixture was further incubated for 30 minutes at 40°C. Tagmented DNA was isolated using 2x SPRI Beads (Beckman Coulter) and amplified through PCR. Fragments smaller than 600 bp were isolated via negative size selection using 0.65x SPRI Beads (Beckman Coulter) and purified using 1.8x SPRI Beads (Beckman Coulter). Quality control was conducted using a a 6000 bioanalyzer (Agilent) in combination with a Qubit 2.0 Fluorometer (Thermo Fisher Scientific). The libraries were subsequently multiplexed in an equimolar pool and sequenced on a NextSeq 500/550 Platform (Illumina), generating at least 30 million reads per sample.

After overall quality control with FastQC and adaptor removal with TrimGalore (v 0.6.10), paired-end reads, each spanning 51 base pairs, were subjected to alignment against the mm10 mouse genome index with Bowtie2 aligner (v 2.5.1) with default parameters, except for the utilization of the “very-sensitive 1000” option. To ensure the quality of aligned reads, those with a quality score below 30 were discarded using Samtools 1.16.1. Mitochondrial reads were excluded from further analysis followed by removal of duplicated reads with Picard (v 2.23.8). The alignment of reads was adjusted to account for TN5 transposase insertions with “alignmentSieve” function from deepTools (v 3.5.1). Peaks were identified using MACS2 (v 2.2.8) with stringent false discovery rate (FDR) threshold of less than 0.01. Regions designated as blacklisted were excised from the analysis (blacklist v 2). Further peak selection for differential accessibility (DA) analysis was carried out by considering peaks that were detected in a minimum of two of the three replicates for each experimental condition and the union of selected peaks for both conditions was generated by GenomicRanges package in the R environment. DA regions were quantified with the csaw package using MACS2 parameters followed by edgeR with trimmed man of M-values (TMM) normalization for DA analysis and significance level was set at 0.5 log-fold change and an FDR < 0.1 with batch correction for samples 2 (CTRL2 and PALM2). Peak annotation was done by ChIPSeeker with up- and downstream regions extending 1 kilobase (kb) from the transcription start site (TSS) based on TxDb.Mmusculus.UCSC.mm10.knownGene database. Heatmaps depicting coverage density were generated utilizing deepTools, focusing on regions extending 1 kb upstream and downstream from the center of each peak. Motif enrichment was performed with JASPAR2022, TFBSTools, MotifDb and chromVAR R packages from bioconductor.

### Histones enrichment

1-2 million cells were collected and resuspended in 0.5 ml of Buffer A (PBS, 0.1% Triton, protease inhibitors, 5 mM NaButyrate). The cells were then pipetted up and down through a 200 μl pipette tip multiple times. Subsequently, the cell suspension was centrifuged for 10 minutes at 5,000 rpm and 4°C. The pellet obtained was resuspended in 50-100 μl of Buffer A+0.1% SDS and sonicated with 10 cycles of 30 seconds on and 30 seconds off to disrupt the chromatin structure.

### Immunoblots

The protein concentration was determined using the bicinchoninic acid assay (BCA, Thermo Fisher Scientific, Waltham, MA, USA). For signaling and mitochondria analysis, 20-30 μg of proteins were loaded, while for histone analysis, 5-10 μg of proteins were loaded on a polyacrylamide gel. Total cell lysates and nuclear fractions were separated using 7-15% polyacrylamide gel and then transferred to nitrocellulose membranes. The membranes were subsequently probed with the listed antibodies. To visualize the results, appropriate secondary antibodies were applied, and the blots were revealed using peroxide solution.

### Histone Digestion and LC/MS analysis

About 5 μg of histones per run per sample were mixed with an equal amount of heavy-isotope labelled histones, which were used as an internal standard for PTM quantification (super-SILAC mix (*83*)) and were separated on a 17% SDS-PAGE gel. A band corresponding to the histone octamer (H3, H4, H2A, H2B) was excised for in-gel digestion with a protocol involving chemical acylation of lysines with propionic anhydride and derivatization of peptide N-termini with phenyl isocyanate (PIC) after trypsin digestion, as described in (*84*). Peptide mixtures were separated by reversed-phase chromatography on an EASY-Spray column (Thermo Fisher Scientific), 25-cm long (inner diameter 75 µm, PepMap C18, 2 µm particles), which was connected online to a Q Exactive HF instrument (Thermo Fisher Scientific) through an EASY-Spray™ Ion Source (Thermo Fisher Scientific). The chromatographic solvents consisted of 0.1% formic acid (FA) in ddH2O (solvent A) and 80% acetonitrile (ACN) plus 0.1% FA (solvent B). Peptides were injected in an aqueous 1% trifluoroacetic acid (TFA) solution at a flow rate of 500 nl/min and were separated with a 50-min linear gradient of 10–45% of solvent B, at a flow rate of 250 nl/min. The Q Exactive instrument was operated in the data-dependent acquisition (DDA) mode, switching between full scan MS and MS/MS acquisition. Survey full scan MS spectra (m/z 300-1350) were analyzed in the Orbitrap detector at a resolution of 60,000-70,000 (m/z 200). The 10-12 most intense peptide ions with charge states between 2 and 4 were sequentially isolated and fragmented by higher-energy collisional dissociation (HCD) with a normalized collision energy setting of 28%. The maximum allowed ion accumulation times were 20 ms for full scans and 80 ms for MS/MS, with a target value for MS/MS set at 1 × 10^5^. The dynamic exclusion time was set to 10 seconds, and standard mass spectrometric conditions for all experiments included a spray voltage of 1.8 kV and no sheath and auxiliary gas flow. The acquired RAW data were analyzed using Epiprofile 2.0 (*85*), selecting the SILAC option, followed by manual validation (*84*). For each histone modified peptide, the % relative abundance (%RA) for the sample (light channel (L)) or the internal standard (heavy channel (H)) was estimated by dividing the area under the curve of each modified peptide for the sum of the areas corresponding to all the observed forms of that peptide and multiplying by 100. Light/Heavy (L/H) ratios of %RA were then calculated and are reported in Dataset S1. The mass spectrometry data have been deposited to the ProteomeXchange Consortium (*86*) via the PRIDE partner repository with the dataset identifier PXD046502.

### Acetyl Co-A assay

CD8+ T cells were cultured for 48 hours in the presence or absence of PA. Afterward, the cells were collected and washed with PBS to enable the analysis of both total and cytoplasmic AcCoA levels.To extract cytoplasmic AcCoA, 1×10^6 T cell samples were homogenized in a solution containing 1% Triton X-100, 20 mM Tris-HCl (pH=7.4), and 150 mM NaCl, while keeping the samples on ice for 10 minutes. Subsequently, the homogenates were centrifuged at 20,000 g for 10 minutes at 4°C to separate the nuclei, and the supernatant obtained after centrifugation was used to measure the AcCoA nucleocytosolic fraction. For quantification of total cellular AcCoA, 1×10^6 T cells were washed with PBS, and the resulting pellet was subjected to two freezing-thawing cycles using liquid nitrogen and either 80% methanol or 5% sulfo-salicylic acid with 50 mM DTT. After that, the samples were deproteinized using a 10 kDa MWCO spin filter prior to conducting the assay. The quantification of AcCoA was performed using the Acetyl-Coenzyme A Assay Kit (Sigma), following the instructions provided by the manufacturer.

### Metabolic assays

Seahorse experiments were conducted on CD8+ T cells cultured for 48 hours with or without PA, using the XF Cell Mito Stress kit from Seahorse Bioscience. The measurements of Oxygen Consumption Rate (OCR) and Extracellular Acidification Rate (ECAR) were performed using XF96 Extracellular Flux Analyzers (Seahorse Bioscience), following the previously described protocols (*87*). In brief, the cells were seeded onto poly-D-lysine-coated 96-well polystyrene Seahorse plates at a density of 200,000 T cells per well and incubated at 37°C for 1 hour for equilibration. The analysis of OCR and ECAR was conducted under basal conditions, and then various compounds were added to induce specific metabolic changes. For the mito stress test, the following compounds were added: oligomycin (1 μM), carbonyl cyanide-4-phenylhydrazone (1.5 μM), and antimycin A/rotenone (1 μM/0.1 μM).

### Metabolomics and lipidomic assays

Liquid chromatography–mass spectrometry (LC-MS) lipidomic analyses were conducted on CD8+ T cells following 48 hours of culture with and without PA. The analysis utilized reverse-phase liquid chromatography (RPLC) coupled with positive- and negative-mode electrospray ionization high-resolution MS. Lipids from the samples were extracted using the Folch extraction method. In brief, cell pellets were resuspended in 100 μl of ammonium formate buffer (50 mM, pH 6.8), and then 400 μl of ice-cold methanol was added. After vortexing and sonication with a sonication probe for 10 pulses at 30% power while cooling on ice, 800 μl of chloroform and 100 μl of water were added to each sample to achieve a final ratio of 4:2:1 chloroform:methanol:water. The samples were vortexed, incubated on ice for 10 minutes, and then centrifuged at 1,000 rpm for 10 minutes. The organic layer containing the lipids was transferred to a clean Eppendorf tube, and the aqueous layer extraction with chloroform was repeated once. The organic layers (lipid layers) were combined and dried down under vacuum.Before MS analysis, the dried lipids were reconstituted in chloroform:methanol (50:50, vol/vol) containing an internal heavy-labeled lipid standard (Splash Lipidomix Mass Spec Standard; Avanti Polar Lipids). The samples were centrifuged and volume adjusted to ensure normalization to the cell count per volume (ml). Global untargeted lipidomic analyses were performed using full MS and data-dependent acquisition analyses on a Q-Exactive HF hybrid quadrupole-Orbitrap mass spectrometer (Thermo Fisher Scientific). RPLC was performed on a Hypersil Gold 1.9-mm, 2.1 x 100-mm column (Thermo Fisher Scientific) held at 40°C. The separation was conducted at 250 μl/min using solvent A (0.1% formic acid in water) and solvent B (0.1% formic acid in isopropanol:acetonitrile:water [60:36:4]) with a specific gradient. The raw data from ultra-performance liquid chromatography/tandem MS were imported, processed, normalized, and reviewed using Progenesis QI v.2.1 (Non-linear Dynamics). Statistical analysis including PCA and P values generation using ANOVA or pairwise comparison was performed to identify statistically significant changes. Tentative annotations were made using accurate mass measurements, isotope distribution similarity, and assessment of fragmentation spectrum matching from the Lipid Maps database. The features were considered differentially expressed if they met both criteria of fold change |R| ≥ 2 and significance (P < 0.05).

### Adoptive cell transfer (ACT) experiments

In vivo adoptive Cell Transfer (ACT) experiments were conducted using the B16-OVA melanoma cell line with antigen OT-I CTL (*88*). The cell line was cultured in RPMI with 10% FBS, 1% PenStrep, and 0.4mg G418 for up to 5 passages before injection. C57B6/J mice were subcutaneously injected with 2×10^5^ cells, and daily monitoring of tumor growth was performed. 10 days post-injection, when tumors reached a measurable size, ACT was administered intravenously using 5×10^6^ OT-I cells previously activated in the presence or absence of PA and SPHK2i. Tumors were assessed and measured every 2 days, and a growth curve was constructed using the formula: Tumor volume = length x width2/2, where length corresponds to the largest tumor diameter, and width represents the perpendicular tumor diameter.

### Analysis of survival and immunotherapy data

Survival analysis was performed on each TCGA project with more than 75 samples (29 datasets). For each project, samples were stratified by lower vs higher than median SPHK2 expression. Survival analysis was performed using the Kaplan-Meier estimator, whose significance was tested using the log-rank test.

Datasets for lung, kidney and skin cancer treated with anti-PD1 drugs were downloaded from the Tumor Immunotherapy Gene Expression Resource database (TIGER) and have the following ids: Braun_2020, GSE78220, GSE91061, phs000452, NSCLC_GSE135222 (*55*, *89–92*). The association between SPHK2 expression and anti-PD1 response was tested using Chi-squared test.

### Quantification and statistical analysis

Flow cytometry data were analyzed using FlowJo v10.2 software. Quantification of Western blots was performed using ImageJ software. Statistical analysis was carried out using GraphPad Prism. The results are presented as mean ± standard error (SEM). Paired t-test was utilized to compare PA-CTL and CTR-CTL, unless otherwise specified. Normal distribution was assessed using the Shapiro-Wilk test.

Metabolomics data were normalized to cell count and/or protein content. From a total of 198 metabolites profiled, 14 metabolites were removed due to the high number of missing values (>20%) and metabolites with less than 20% missing values were imputed using the median approach. Due to the high dimensionality and collinear nature of the data, logistic regression with elastic net penalty and a binomial modality (alpha = 0.5) was implemented in the “glmnet” (R package, version 3.0-2) to select the metabolites that were associated with the PA exposure, as compared to the CTRL. Biological significance was considered for those metabolites with a regression coefficient ≥ |0.2|. A pathway score was then created for each of the significant metabolites and graphically represented with a bubble plot (R package, “ggplot2”, version 3.4.3). Lipidomic analysis was conducted on the lipid compounds obtained by the negative acquisition whose annotations were supported by a precursor mass and MS/MS. Thus, we adopted the analytical pipeline of “lipidr” R package (version 2.10) for a total of 114 lipids. The initial step of this workflow included the data inspection and integration with reference information about lipid class, total chain length and unsaturation levels. Because of missing information in the annotated lipids we were able to only parse 107 compounds. We therefore run a differential abundances analysis, based on a moderated T-test, as part of the analytical pipeline. 58 metabolites were significantly associated to PA exposure and a compositional heatmap of these metabolites was created with “ComplexHeatmap” R package (version 2.12.1)). Histone PTM L/H ratios in PA-treated cells were compared to the corresponding control cells by paired Student’s T test.

All graphs display mean ± SEM, and statistical significance was represented by **** p-value < 0.0001, *** p-value < 0.001, ** p-value < 0.01, and * p-value < 0.05.

---

**Figure S1.**
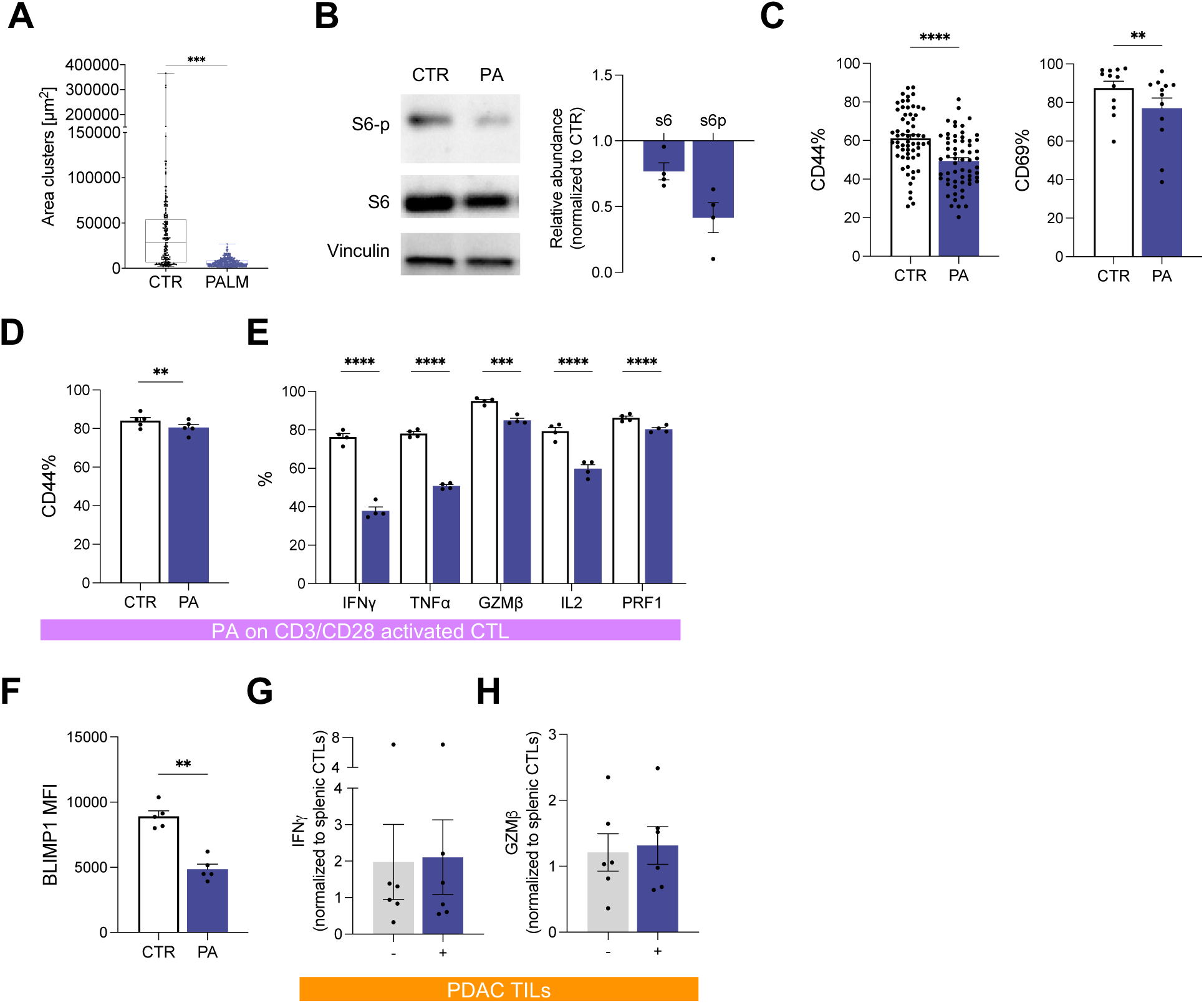
(relative to Figure 1). **(A).** Quantification of clusters area after activation in CTR and PA-CTL (N=3). **(B).** Western blot representative image and relative abundance of S6 and S6-p in PA-CTL normalized on CTR-CTL (N=4). Vinculin is shown as a loading control. **(C).** Quantification of CD44 (N=61) and CD69 expression (N=12). **(D-E).** Effect of PA exposure on CD3/CD28 activated CTL. Quantification of CD44 (**D,** N=4) and cytokines production (**E,** N=4). **(F).** BLIMP1 expression (N=5). **(G-H).** Quantification of GZMB (**G**) and IFNγ(**H**) production from TILs isolated from PDA mouse model with (+) and without (-) restimulation with PMA-ionomycin. (N=6)

**Figure S2.**
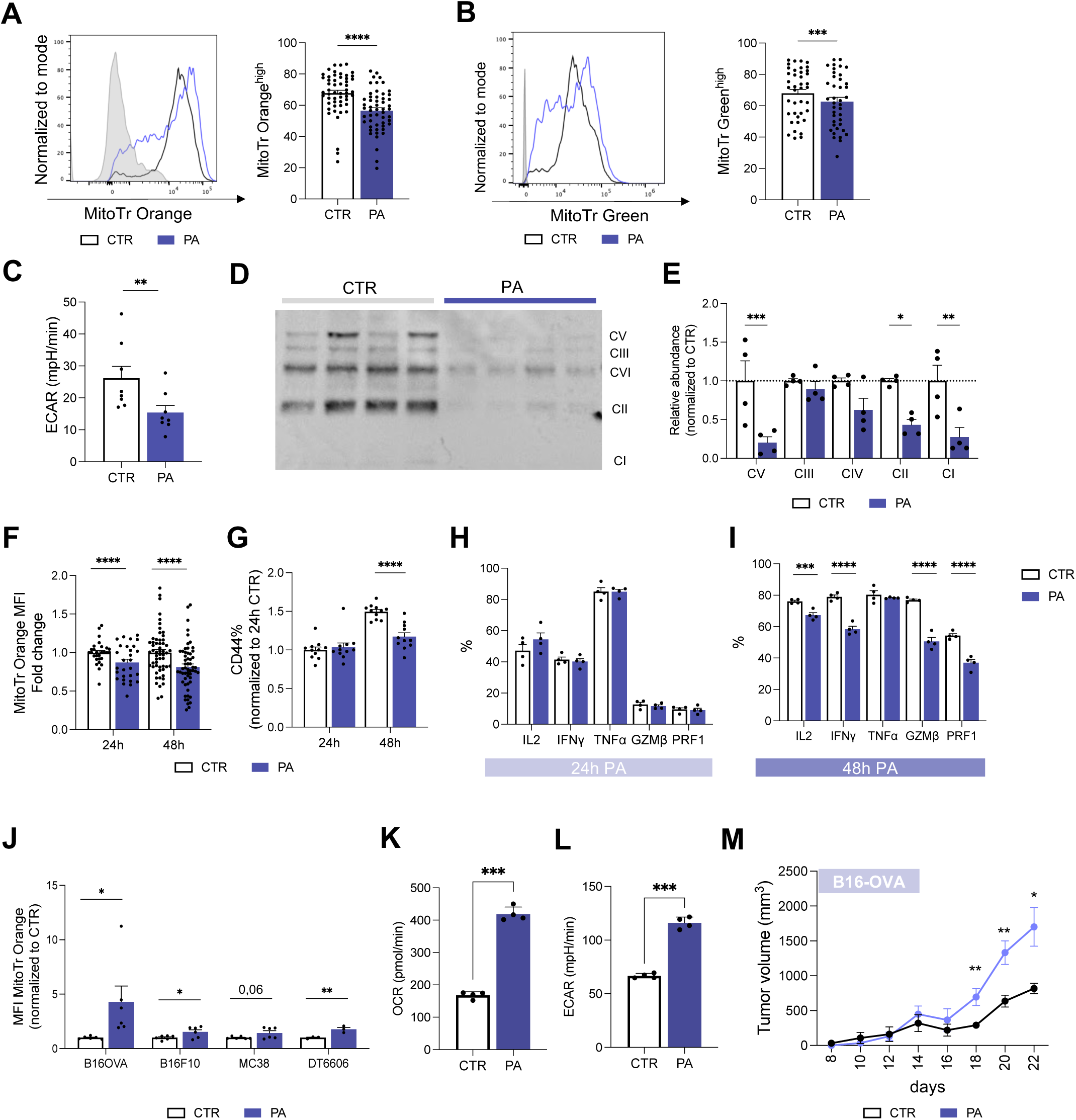
(relative to Figure 2). **(A-B).** Mitochondrial potential **(A,** N=60) and mass **(B,** N=44) based on flow cytometry. **(C-D).** Western blot of electron transport chain complexes (**C**) and relative quantification (**D**) (N=6). **(E).** Quantification of basal Extracellular Acidification Rate (ECAR, N=8). **(F-I).** Quantification of mitochondrial activity (**F**), CD44 expression (**G**) and effector cytokines production (**H-I**) after 24h or 48h of PA treatment (N=4/28). **(J-L).** Characterization of cell lines treated with PA. Mitochondrial potential (**J,** N=6), metabolic profile measured as basal OCR (**K,** N=3) and ECAR (**L,** N=3) and tumor growth curves (**M,** N=6 per group) of tumor cell lines ± PA (N=6).

**Figure S3.**
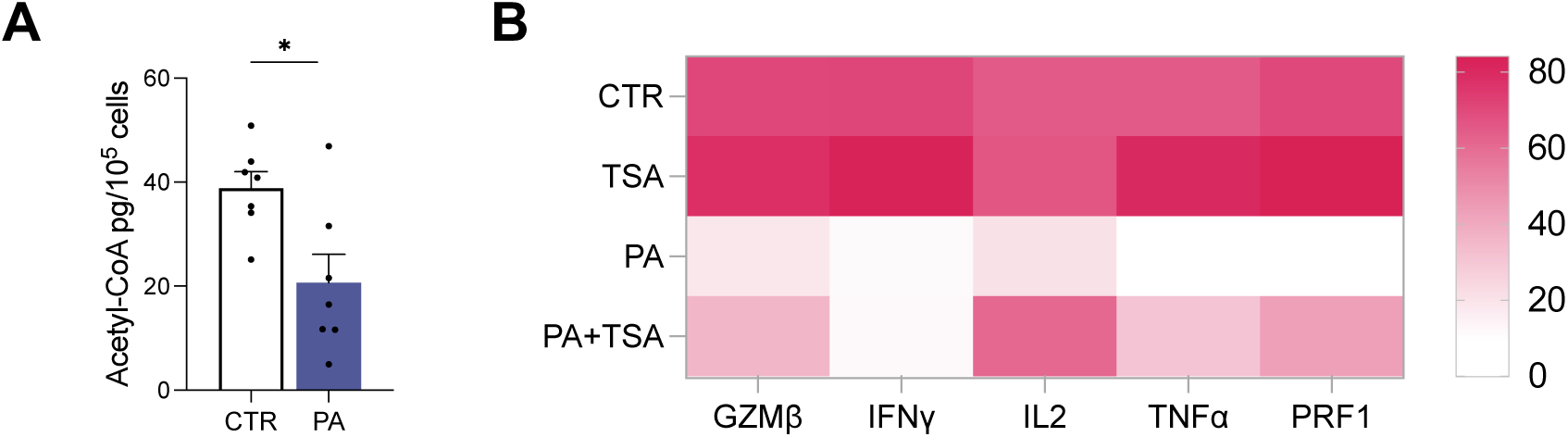
(relative to Figure 3). **(A).** Acetyl-CoA quantification in CTL±PA **(B).** Cytokines production after 48h of culture with and without PA and with and without 12.5nM TSA.

**Figure S4.**
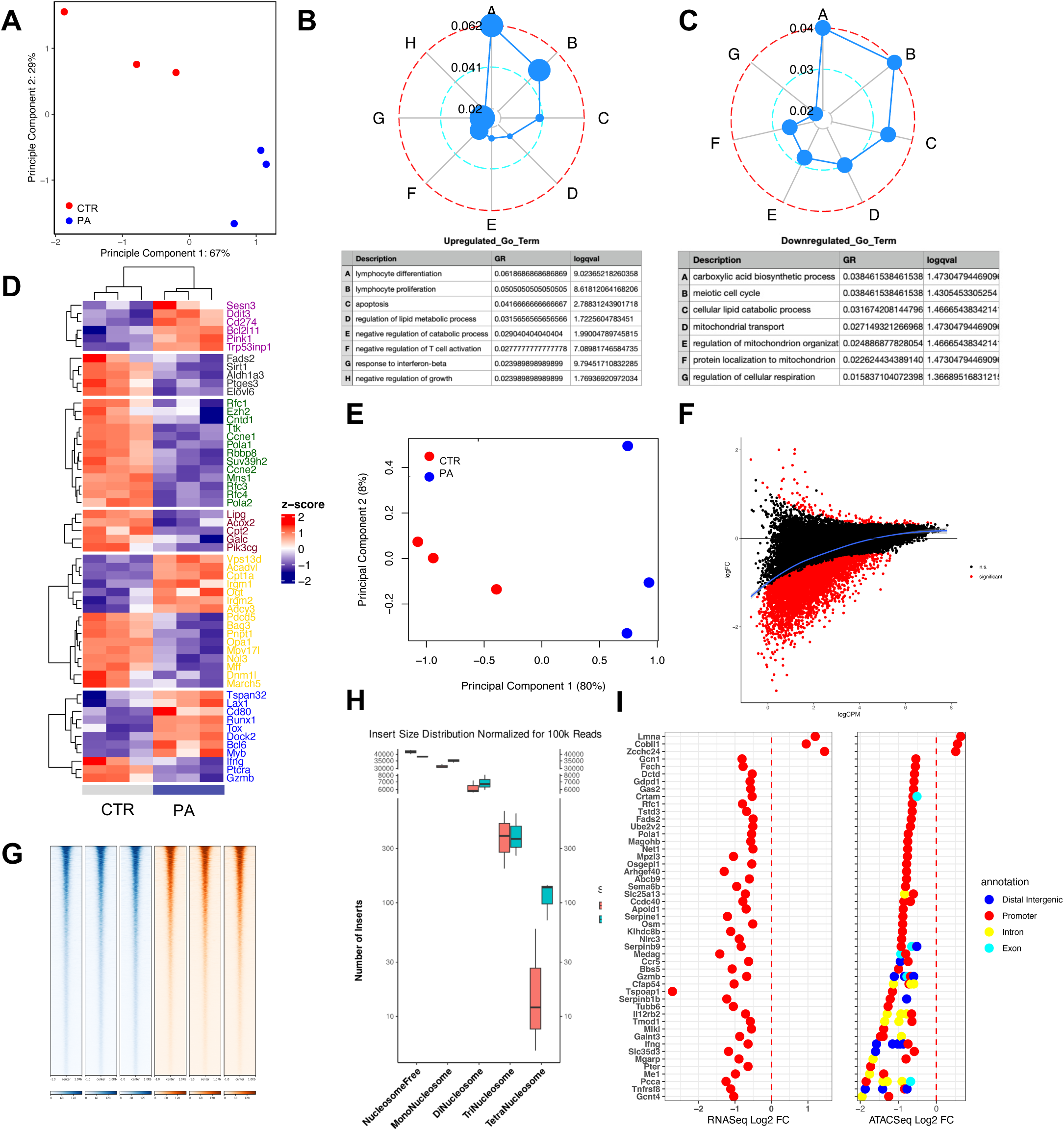
(related to Figure 4). (**A**) PCA plot for RNASeq based on rlog values plotted with ggplot. (**B,C**) Radar plots depicting 8 GO Term enrichment from genes upregulated and 7 GO Terms enriched by genes downregulated by palmitate treatment in RNASeq. Concentric circles show gene ratio as calculated by cluster profiler and dot sizes the negative log10 of the q-value. (**D**) Heatmap showing the expression of a number of manually selected genes representing the pathways and functions color-coded as shown in the legend. Rlog values were normalized row-wise and clustering was done within each group. The heatmap was generated with ComplexHeatMap package. (**E**) PCA plot for ATAC-Seq data generated with plotMDS function of edgeR package. (**F**) MA plot of DA analysis of ATACSeq data. PA-treated cells are on the positive y-axis and control on the negative y-axis with loess curve fit. (**G**) Heatmaps and average plots showing ATACSeq signals for each sample for 1kb up- and downstream of peak center. (**H**) Insert size quantification for chromatin accessibility based on log-insert sizes plot normalized for 100000 reads. (**I**) DotPlots of commonly up- and downregulated genes in RNASeq and ATACSeq. Y-axis shows gene names and x-axis the log-fold change between palmitate-control. For ATACseq, genomic annotation of each peak is color-coded a shown in the legend.

**Figure S5.**
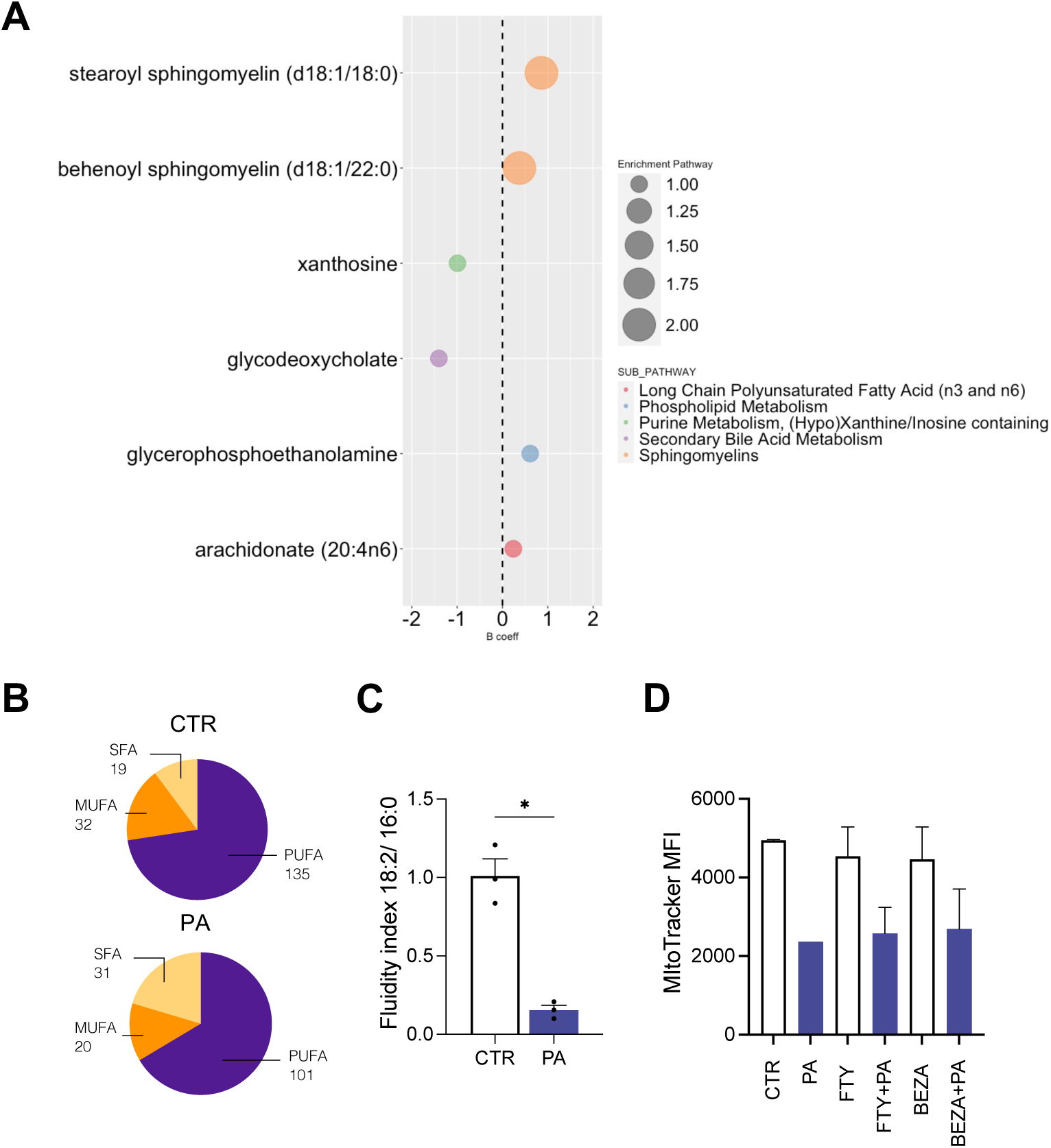
(relative to Figure 5). **(A).** Enrichment analysis of metabolites positively associated with PA treatment in CTL (N=8) **(B).** Quantification of saturated (SFA), mono-unsaturated (MUFA) and poli-unsaturated fatty acids (PUFA) fatty acids in CTL±PA (N=3). **(C).** Fluidity index (18:2/ 16:0) of CTL±PA (N=3)**. (D)**. Mitochondrial potential in CTL±PA alone or in combination with FTY (Fingolimod, 200nM) or Beza (Bezafibrate, 100uM) (N=3).

**Figure S6.**
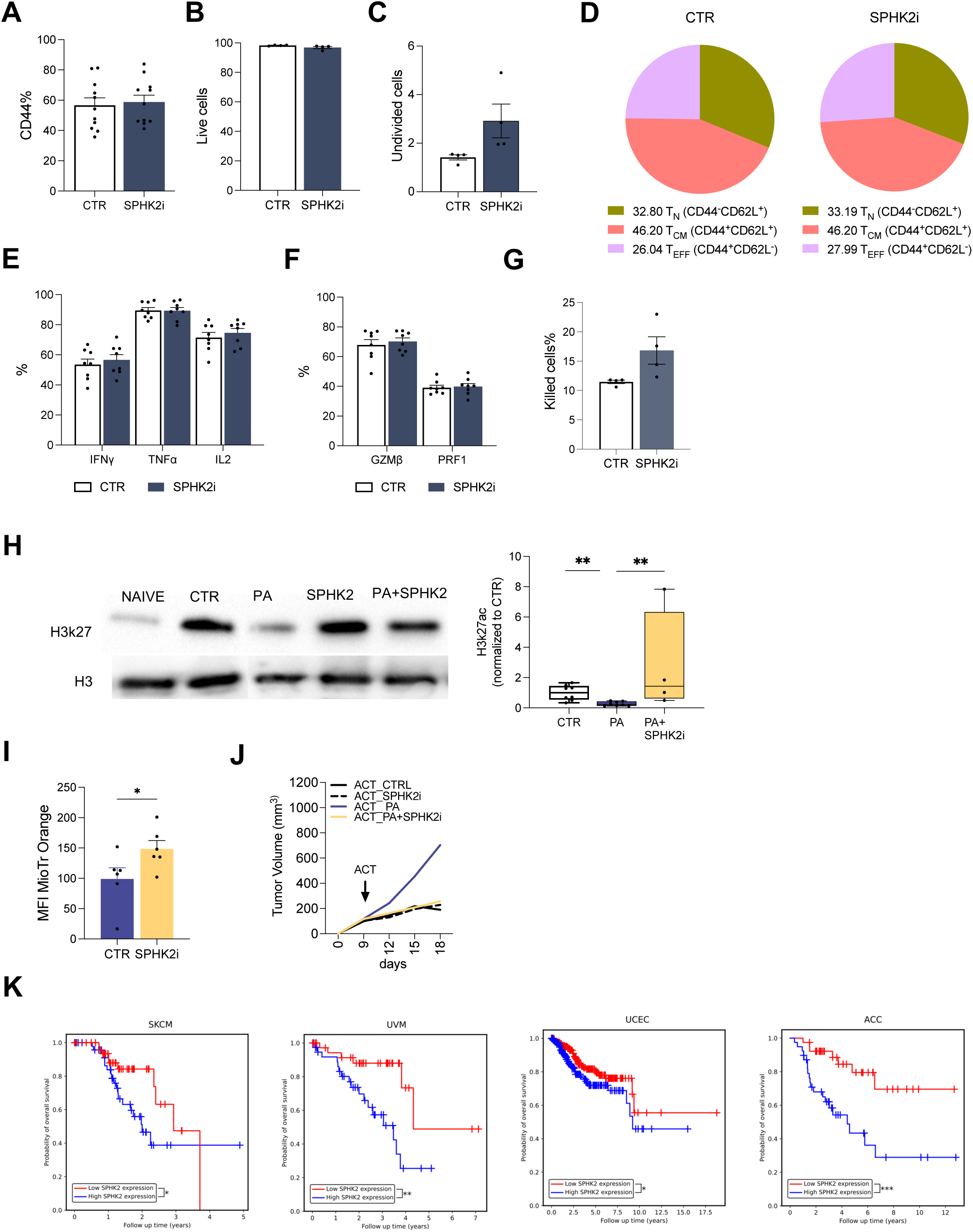
(related to Figure 6). **(A-G).** Effect of SPHK2i on CTL activation and function measured as CD44 expression (**A**) viability (**B**) proliferation **(C),** differentiation profile **(D),** cytokines and effector molecules production **(E-F)** and percentages of killed B16OVA tumor cells after coculture with OTI ±SPHK2i in 1:1 ratio. (**G,** N=4). **(H).** Western blot and relative quantification of H3k27ac with and without SPHK2i in PA-CTL and CTR-CTL **(I).** Quantification of mitochondrial potential in tumor infiltrating lymphocytes (TIL) isolated from PDAC models after 24h culture ± SPHK2i (N=6). **(J).** Growth curves of B16OVA tumor bearing mice treated with CLT ± PA±SPHK2i (N=4/15)**. (K).** Survival plots of low vs high SPHK2 expression samples for SKCM (p=0.0471), ACC (p=0.0008), UCEC (p=0.0250), and UVM (p=0.0046) TCGA projects.

